# Transcytosis maintains CFTR apical polarity in the face of constitutive and mutation-induced basolateral missorting

**DOI:** 10.1101/411322

**Authors:** Aurélien Bidaud-Meynard, Florian Bossard, Andrea Schnúr, Ryosuke Fukuda, Guido Veit, Haijin Xu, Gergely L. Lukacs

**Affiliations:** Department of Physiology, McGill University, Montréal, QC, H3G 1Y6, Canada; Department of Biochemistry, McGill University, Montréal, QC, H3G 1Y6, Canada

## Abstract

Apical polarity of cystic fibrosis transmembrane conductance regulator (CFTR) is essential for solute and water transport in secretory epithelia and can be impaired in human diseases. Maintenance of apical polarity in the face of CFTR non-polarized delivery and compromised apical retention of mutant CFTRs lacking PDZ-domain protein (NHERF1) interaction, remains enigmatic. Here we show that basolateral CFTR delivery originates from biosynthetic (~35%) and endocytic (~65%) recycling missorting. Basolateral channels are retrieved via basolateral-to-apical transcytosis, enhancing CFTR apical expression by two-fold and suppressing its degradation. CFTR transcytosis is microtubule-dependent but independent of Myo5B-, Rab11- and NHERF1 binding to its C-terminal DTRL motif in airway epithelia. Increased basolateral delivery due to compromised apical recycling and accelerated internalization upon impaired NHERF1-CFTR association is largely counterbalanced by CFTR efficient basolateral internalization and apical transcytosis. Thus, transcytosis represents a previously unrecognized but indispensable mechanism for maintaining CFTR apical polarity by attenuating its constitutive and mutation-induced basolateral missorting.

## INTRODUCTION

The anion selective channel CFTR, a member (ABCC7) of the ABC transporter superfamily, is predominantly expressed at the apical plasma membrane (PM) of secretory and resorptive epithelia (Riordan, 2008). CFTR maintains the ionic, composition and volume homeostasis of the airway surface liquid, prerequisites for the physiological regulation of mucocilliary clearance and innate immune response of airway epithelia, and to prevent uncontrolled infection and inflammation of the lung, the primary causes of morbidity and mortality in cystic fibrosis (CF) (Cutting, 2015; Rowe et al., 2005). Inherited funtional expression defect of CFTR at the apical PM of epithelia causes CF, while acquired loss of CFTR expression in airway epithelia and pancreatic duct is associated with chronic obstructive pulmonary disease (COPD), asthma and pancretitis, respectively (Schnur et al., 2016).

Four vesicular transport routes determine CFTR steady-state expression at the PM: biosynthetic secretion, slow constitutive internalization and recycling, as well as endo-lysosomal/autophagosomal degradation (Ameen et al., 2007; Fu and Sztul, 2009; Gentzsch et al., 2004; Sharma et al., 2004). Given the long cellular and PM half-life of CFTR (Okiyoneda et al., 2018; Varga et al., 2008), slow internalization rate with high fidelity of apical recycling is essential to maintain Cl^-^ transport capacity by minimizing the channel premature degradation and basolateral missorting.

In polarized epithelia, apical and basolateral PM proteins partition between apical sorting (ASE), apical recycling (ARE), basolateral sorting (BSE), and common recycling (CRE) endosomes, respectively (Stoops and Caplan, 2014). Previous studies have shown that apically internalized CFTR is confined to EEA1^+^ ASE and recycles via Rab11^+^ ARE in airway cells (Cholon et al., 2009; Swiatecka-Urban et al., 2007), in a PKA-dependent manner (Holleran et al., 2013), but the molecular basis of rapid degradation of basolaterally delivered and natively folded CFTR is not known. Nevertheless, forced missorting of CFTR to the basolateral PM suppressed the transepithelial anion secretion efficiency (Farmen et al., 2005).

Apical confinement of PM proteins can be achieved either by direct apical targeting from the *trans*-Golgi network (TGN) (Takeda et al., 2003), indirectly by transcytosis from the basolateral PM (Anderson et al., 2005) or by non-polarized delivery with selective retention at the apical PM and rapid degradation from the basolateral PM, as described for CFTR (Swiatecka-Urban et al., 2002). The association of PDZ (Postsynaptic density/Disc large/ZO-1) domain adaptor proteins with PM proteins exposing a PDZ binding motif can serve as polarized targeting and/or retention signal at both the apical and basolateral PM (Brone and Eggermont, 2005). NHERF1/2 binding to CFTR C-terminal PDZ binding motif (DTRL^1480^) has been implicated in the channel apical targeting, retention, confined lateral diffusion and endocytic recycling (Haggie et al., 2004; Moyer et al., 1999; Swiatecka-Urban et al., 2002) by tethering it via ezrin to the subapical actin network (Short et al., 1998; Sun et al., 2000). Deletion of the DTRL motif caused by the S1455X premature termination codon, however, manifested in the isolated elevation of sweat Cl^-^ concentration in the absence of lung and pancreatic phenotype in the background of an in frame deletion of exon 14a, which encodes a non-functional CFTR (Mickle et al., 1998). These observations with extensive mutagenesis studies (Benharouga et al., 2003; Milewski et al., 2005; Milewski et al., 2001; Ostedgaard et al., 2003) suggested that CFTR tethering to the subapical cytoskeleton via NHERF is not essential for the channel apical polarity. To identify additional mechanism(s) that preserve CFTR polarity in the absence of NHERF tethering function, we investigated the candidate trafficking pathways, sorting signals, and molecular players that may be involved in CFTR polarity development.

Here we show that basolateral-to-apical (or apical) transcytosis is indispensable to maintain CFTR apical polarity and metabolic stability in the face of limited fidelity of the channel apical biosynthetic secretion and recycling from apical endosomes, which both induce its basolateral missorting. The microtubule-dependent, but Myo5B-independent CFTR transcytosis in airway cells avoids both apical and basolateral endocytic recycling compartments but traverses EEA1^+^ sorting endosomes. Furthermore, apical transcytosis counteracts the augmented basolateral missorting of C-terminal truncated CFTR, providing a plausible explanation for the isolated resorbtive defect in the sweat gland of individuals harboring S1445X-CFTR.

## RESULTS

### Human airway epithelial models for studying CFTR polarized sorting

To investigate polarized trafficking, CFTR containing a 3HA-tag in its 4th extracellular loop was expressed in CF <u>b</u>ronchial <u>e</u>pithelial cells (CFBE41o- or CFBE) under the control of the tetracycline transactivator (Ehrhardt et al., 2006; Veit et al., 2012). Filter-grown CFBE cells were cultured for 3-5 days post-confluence and CFTR expression was induced with doxycycline (+dox) to reach similar or lower expression to that in Calu-3 cells, a lung adenocarcinoma epithelia cell line endogenously expressing CFTR (Figure 1A). The mass difference between the endogenous and tagged CFTR can be attributed to 3HA-tag and altered complex-glycosylation, demonstrated by the mobility shift observed after glycosidase digestion (Figure S1A), but had no significant impact on the channel folding, trafficking, stability and function (Peters et al., 2011). As a second model, CFTR-3HA was expressed in conditionally reprogrammed human primary bronchial epithelial cells (CR-HBE) differentiated under air-liquid interface for ≥4 weeks after lentiviral transduction (Figure 1B). Hallmarks of airway epithelia differentiation were documented: tight-junctional localization of ZO-1, mucin 5 accumulation in goblet cells, accumulation of acetylated tubulin in cilia, as well as development of transepithelial electrical resistance (>500 Ω/cm^2^), airway surface liquid (ASL, ~10μm), and polarized H^+^/K^+^ ATPase and transferrin receptor (TfR) expression (Figures 1C, S1B-D and Methods). Apical localization and function of exogenously expressed CFTR-3HA was demonstrated by immunostaining and the presence of dox-inducible and CFTR inhibitor (Inh_172_)-sensitive PKA-activated short circuit current (Isc) (Figures 1D-E and 6I).

**Figure 1.**
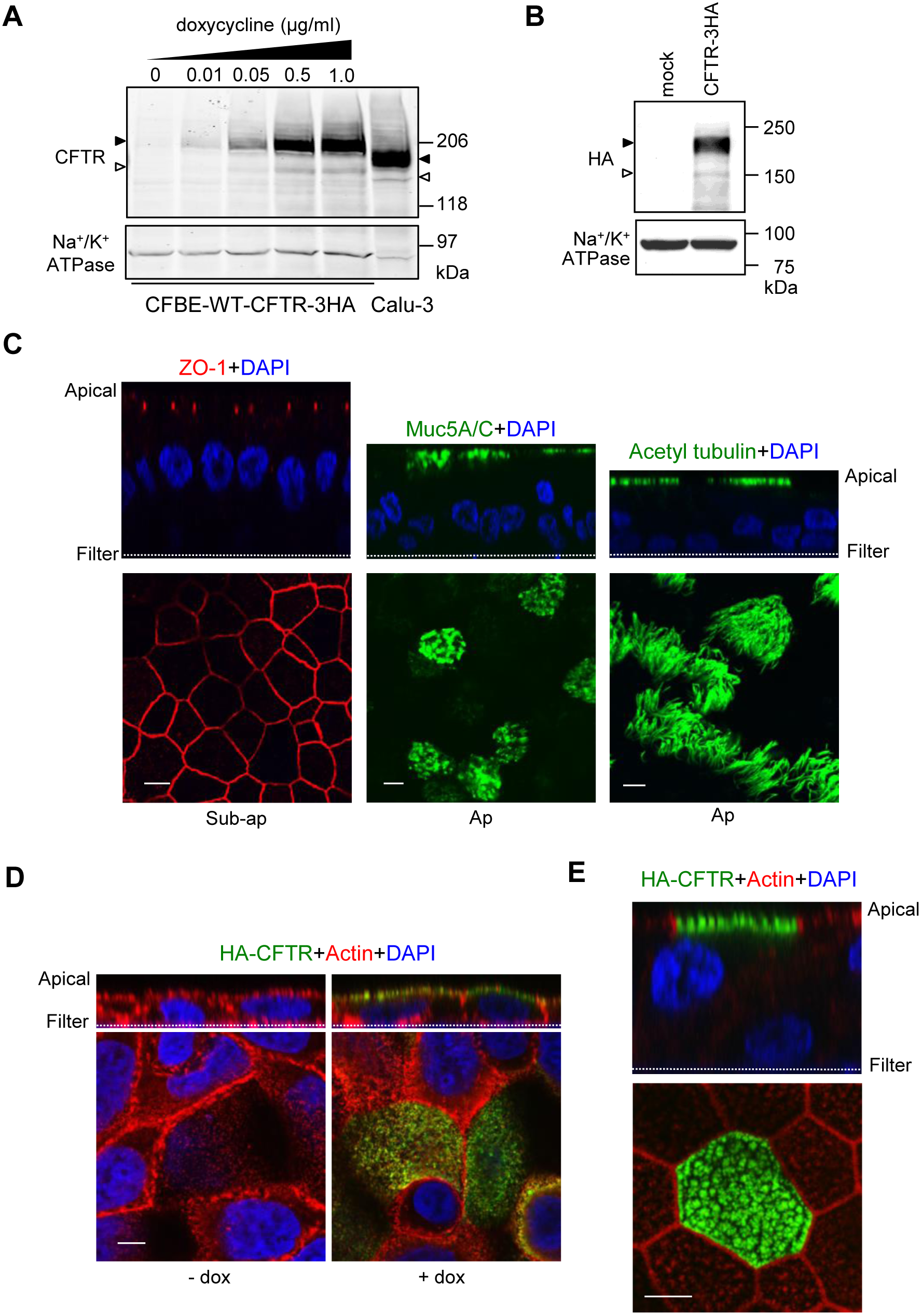
Characterization of human CF bronchial epithelial models to study CFTR polarized sorting. (A-B) Immunoblot analysis of CFTR-3HA expression (A) in CFBE after 3 days doxycycline induction and (B) following lentivirus transduction of CR-HBE. For comparison, equal amount of Calu-3 lysate was loaded (A). Filled and empty arrowheads indicate complex- and core-glycosylated CFTR, respectively. Na^+^/K^+^ ATPase was probed as loading control. (C) CR-HBE were differentiated at air liquid interface (ALI) and stained for tight junction (ZO-1, red), mucin 5 (muc5A/C, green), cilia (acetylated-tubulin, green), and nuclei (DAPI, blue). Cells were visualized by LCFM. Top and lower panels are vertical and horizontal optical sections, respectively. (D-E) WT-CFTR-3HA is apically expressed in CFBE (D) and CR-HBE (E) cells. Cells were stained for WT-CFTR-3HA (green), actin (red) and DAPI (blue). Scale bar: 5 μm. Representative of at least 3 independent experiments.

### CFTR is constitutively transcytosed in human airway epithelia

Newly synthesized CFTR is targeted randomly to both apical and basolateral PM in MDCK cells (Swiatecka-Urban et al., 2002). To study CFTR biosynthetic targeting in CFBE, we determined the kinetics of domain-specific appearance of a cohort of newly translated CFTR, containing an HRP-tag in its 4th extracellular loop (Veit et al., 2014). HRP activity, as a surrogate measure of CFTR PM appearance, was monitored after dox-induction of CFTR-HRP transcription in CFBE and MDCK cells. CFTR-HRP activity appeared simultaneously at the apical and basolateral PM (236 min *vs* 242 min in CFBE; 164 min *vs* 156 min in MDCK, Figure S1E-G), confirming the non-polarized biosynthetic secretion of CFTR in both epithelia (Swiatecka-Urban et al., 2002). Considering that basolateral sorting of CFTR takes place at the TGN, we hypothesized that natively folded channels are targeted for apical transcytosis instead of lysosomal degradation.

To assess apical transcytosis, we monitored the postendocytic fate of basolateral CFTR-3HA after labeling the channel with anti-HA antibody (Ab) capture from the basolateral compartment of filter-grown CFBE and CR-HBE at 37°C for 3h. CFTR-anti-HA Ab complexes were accumulated at the apical PM as well as intracellularly (Figure 2A, *left panel*), visualized by A488-conjugated secondary Ab and laser confocal fluorescence microscopy (LCFM). Following 2h chase, the intracellular CFTR staining was diminished (Figure 2A, *middle panel*). No staining was detected in the absence CFTR (Figure S1H), indicating that the primary Ab staining cannot be attributed to non-specific binding or CFTR-independent Ab transcytosis. We used similar assays to confirm CFTR transcytosis by electron microscopy (EM) with post-embedding secondary Ab decorated nanogold labeling. Anti-HA-CFTR complex accumulation at the apical or basolateral PM was detected following anti-HA labeling from contralateral compartments only in CFTR expressing CFBE (+dox, 3h, 37°C) (Figures 2B-C and S2A), suggesting that CFTR undergoes bidirectional transcytosis.

**Figure 2.**
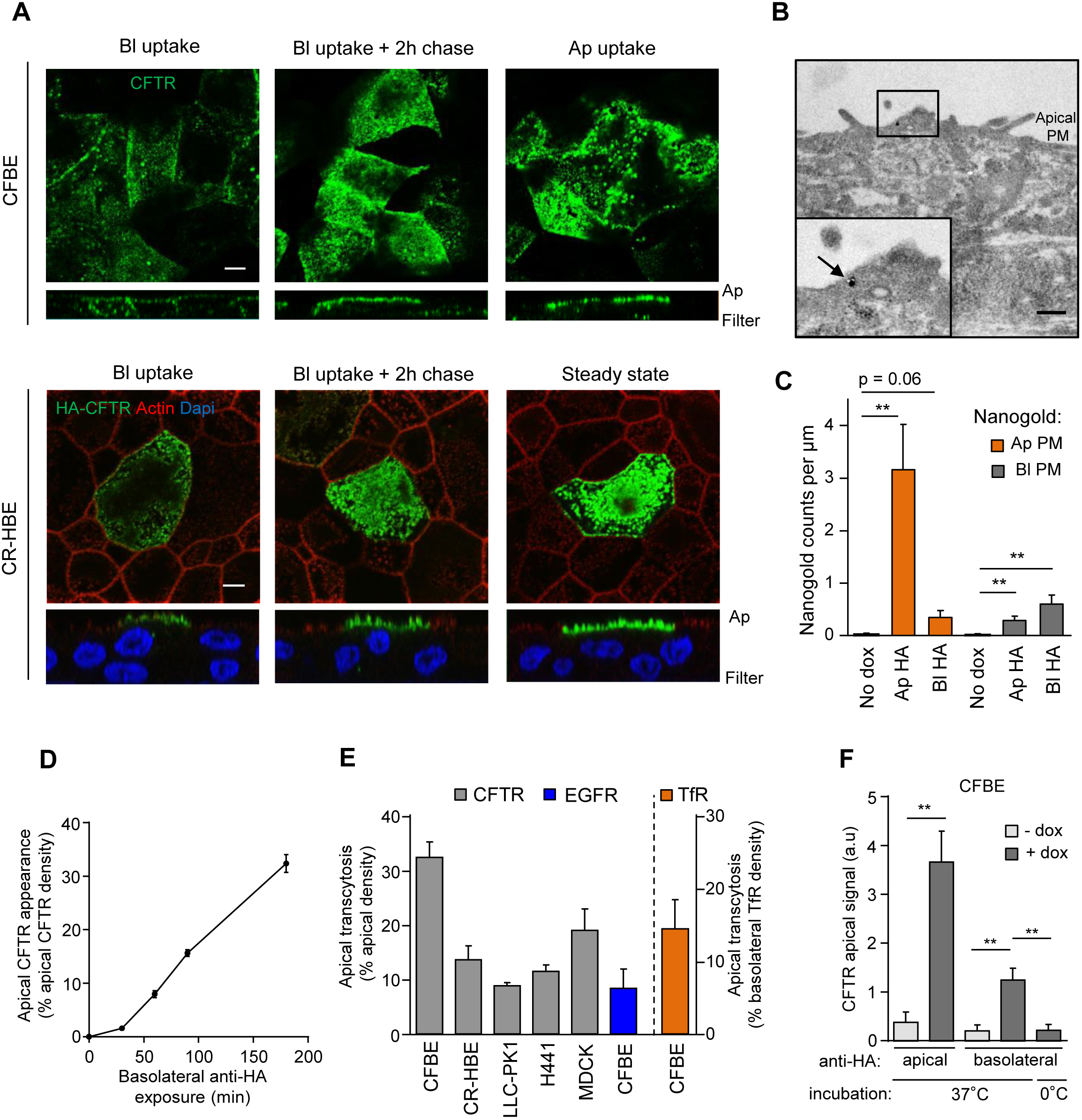
CFTR is apically transcytosed in human airway epithelia. (A) CFTR-3HA was labeled with anti-HA Ab capture (3h, 37°C) from the basolateral (Bl) or the apical (Ap) compartment in CFBE or CR-HBE. Cells were chased for 2h (37°C) in the absence of anti-HA Ab. CFTR-anti-HA Ab complexes were visualized with A488-anti-mouse Ab and LCFM. Lower panels represent z-sections. Scale bar: 5 μm. (B-C) Monitoring CFTR transcytosis with immuno-EM. (B) Immuno-EM detection of CFTR-3HA at the apical PM after basolateral labeling with anti-HA (3h, 37°C) in CFBE. CFTR-anti-HA complexes labeled with secondary Ab decorated nanogold (arrow). Insert represents a higher magnification of the ROI. Scale bar, 0.5 μm. (C) CFTR expressing or non-expressing (no dox) CFBE cells were labeled either by apical (Ap) or basolateral (Bl) anti-HA Ab capture for 3h (37°C). CFTR-anti-HA complexes were visualized with anti-mouse Ab decorated nanogold particle as in panel B. The number of nanogold particle per μm was counted along the apical and basolateral PM segments (n=21-34 ROI/condition in +dox and no-dox, n=2-3 independent experiments, Mann-Whitney U-test). (D) CFTR apical transcytosis was measured over time as in Figure S2B (n=2-10). (E) Apical transcytosis of CFTR-3HA, and endogenous EGFR, and TfR were measured as described in Methods in the indicated cells (n=3-10). (F) Apical transcytosis of CFTR is abolished when the channel is basolaterally labeled by anti-HA Ab for 3h on ice or CFTR transcription was not induced (no dox) (n=4). Data are means ± SEM on each panel, ** p<0.01.

### Determination of CFTR apical transcytotic flux

To estimate the CFTR transcytotic flux, we labeled basolateral channels by anti-HA Ab capture and detected the transcytosed CFTR-Ab complexes by PM-ELISA at the apical PM (Figure S2B, *right*). The amount of transcytosed CFTR amount was expressed as the percentage of the steady-state apical CFTR PM density (Figure S2B, *left*). CFTR accumulation at the apical PM was a function of basolateral Ab-labeling time (0.5-3h) at 37°C (Figure 2D). After 3h of Ab capture, 32 ±3% and 14 ±3% of the apical PM pool was labeled by transcytosis in CFBE and CR-HBE, respectively (Figure 2E). These values, due to constitutive internalization of transcytosed CFTR-anti-HA Ab complexes, are likely underestimated. The channel transcytosis remained unaltered after reducing the dox-induction of CFTR expression to that of ~25% in Calu-3 cells (Figure S2C).

Several evidences indicate the specificity of the ELISA-based transcytosis assay. 1) Transepithelial or paracellular Ab transport was ruled out by the lack of HRP-conjugated Ab translocation from the basolateral into the apical compartment at 37°C (Figure S2D-E). 2) Neither non-specific mouse or rabbit Ab was apically transcytosed, indicating that FcRn- or pIgR-mediated Ab transport cannot contribute to the transcytotic signal (Figure S2F). Likewise, the contribution of fluid phase transcytosis of anti-HA Ab was negligible (Figure S2F, Ap HA vs Ap + Bl HA). 3) CFTR transcytosis was completely abolished on ice or in the absence of CFTR expression (Figure 2F). 4) We showed previously that primary and secondary Abs do not alter CFTR PM turnover (Sharma et al., 2004). 5) We confirmed that endogenously expressed TfR and EGF-receptor (EGFR) are also missorted by apical transcytosis in CFBE (Figure 2E) despite their highly polarized basolateral PM expression. Jointly, these results suggest that CFTR belongs to a handful of PM proteins (e.g. TfR, aquaporin 2 (AQP2) and the Cu-ATPase) whose polarized missorting is compensated by transcytosis.

### Contribution of biosynthetic and endocytic missorting to CFTR basolateral delivery

Basolateral delivery of newly synthesized CFTR may originate from the endoplasmic reticulum by non-conventional trafficking (Gee et al., 2011; Yoo et al., 2002), the TGN (Swiatecka-Urban et al., 2002) and/or from the apical PM via basolateral (apical-to-basolateral) transcytosis. To evaluate the biosynthetic pathway contribution to CFTR basolateral delivery, apical transcytosis was measured following termination of CFTR transcription/translation. After 24h dox washout (dox-OFF), CFTR translation was completely abrogated, confirmed by metabolic pulse-labelling (Figure 3A). Biosynthetic arrest decreased CFTR apical transcytosis by 33.6 ±4.8% in CFBE. CFTR apical transcytosis was similarly reduced (40-50%) in polarized kidney (MDCK, LLC-PK1) and airway (H441) epithelia (Figure 3B).

**Figure 3.**
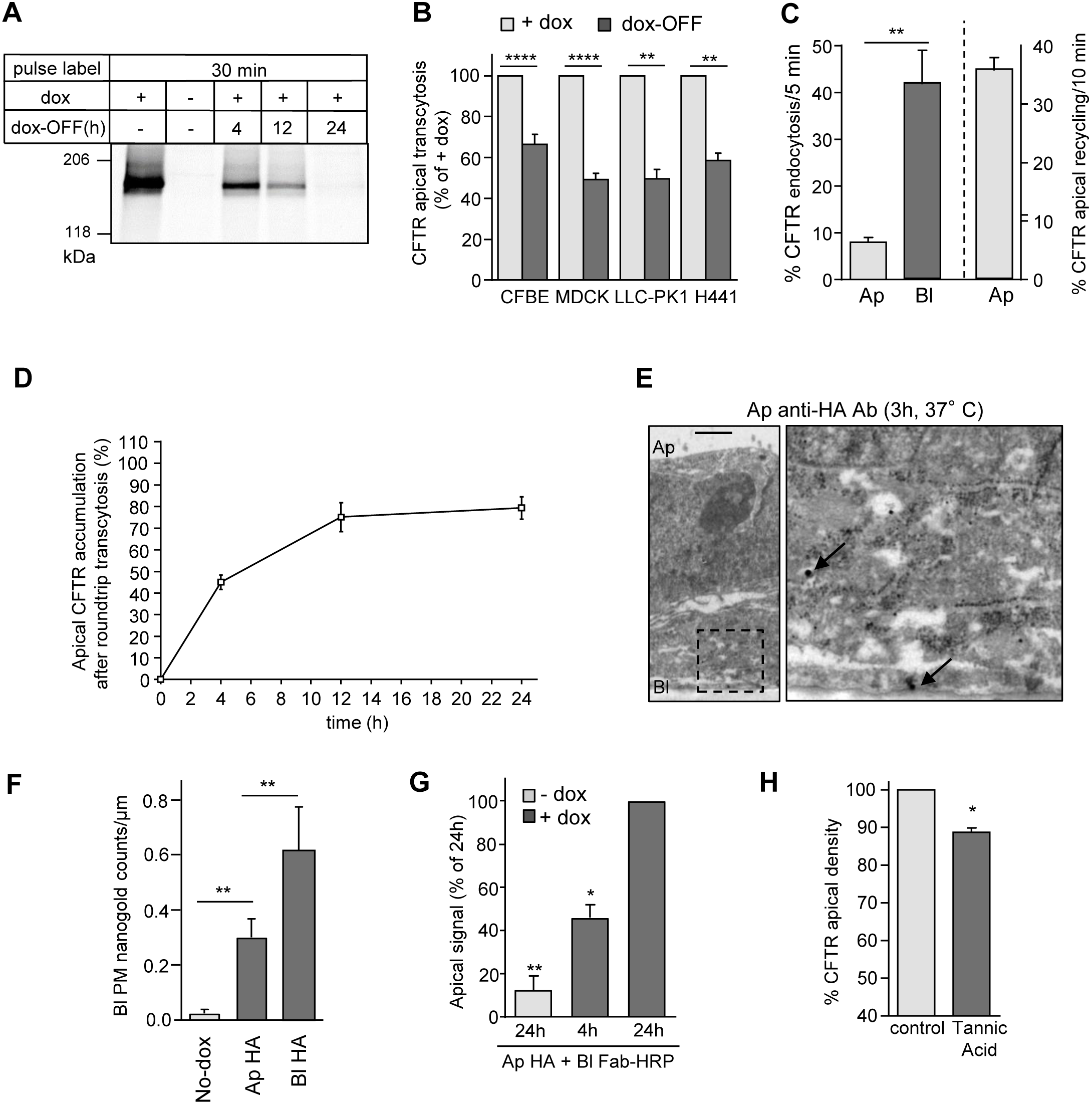
Apical transcytosis retrieves basolaterally missorted CFTR from the biosynthetic and apical recycling pathways. (A) Onset of CFTR translational arrest in dox-OFF CFBE. Translational inhibition was measured by metabolic pulse labeling with ^35^S-metionine/^35^S-cysteine (30min) after 0-24h of dox-OFF by fluorography. (B) CFTR apical transcytosis after 24h dox-OFF in the indicated epithelia (n=3-17). (C) CFTR endocytosis and recycling in CFBE monitored by PM ELISA (n=7-29, two-tailed unpaired t-test). (D) Apical CFTR labeling by basolateral-to-apical transcytosis. CFTR was labeled by continuous basolateral anti-HA capture at 37°C. Apical CFTR-Anti-HA Ab complexes were measured by PM-ELISA (n=4-5). (E-F) CFTR basolateral transcytosis detection by immuno-EM in CFBE after apical capture of anti-HA Ab (3h, 37°C). Right micrograph is a higher magnification of the selected area. Scale bar: 1μm; arrows, CFTR-anti-HA-nanogold-anti-mouse complex. (F) quantification of nanogold particles at the basolateral PM after apical anti-HA Ab uptake (n=21-31 R.O.I.s/condition ROI/condition in +dox and no-dox, n=2-3 independent experiments, Mann-Withney U-test). (G) Round-trip CFTR transcytosis was monitored by simultaneous exposure of CFBE to anti-HA and anti-mouse Fab-HRP in the apical and basolateral compartment, respectively, for 4h or 24h. Transcytosed CFTR was quantified by PM-ELISA (n=3). (H) Inhibition of transcytosis accelerates apical CFTR turnover. After blocking newly synthesized CFTR arrival (dox-OFF, 24h), CFBE was exposed basolaterally to 0.5% tannic acid (15min, 37°C) or mock treatment. Apical CFTR density was measured after 1h incubation at 37°C with PM-ELISA (n=3) and expressed as % of control. Ap: apical; Bl: basolateral. Data are means ± SEM on each panel, n.s. not significant, * p<0.05, ** p<0.01, **** p<0.0001.

Under steady-state condition, we detected only a small fraction of the total CFTR-HRP (2.4 ±0.6%) or CFTR-3HA (3.3 ±0.4%) expression at the basolateral PM by HRP activity and PM ELISA, respectively (Figure S3A). The modest steady-state basolateral density of CFTR can be attributed to the concerted result of the surprisingly rapid basolateral (41.8 ±7.3%/5min) and the >5-fold slower apical (8.0 ±1.0%/5 min) internalization rate, in concert with efficient apical endocytic recycling (35.8 ±2.2%/10min) of CFTR (Figure 3C). These processes are complemented by CFTR constitutive apical transcytosis to preserve the channel polarity, substantiated by the observation that at least 75% of the apical CFTR pool could be labeled by transcytosing CFTR-anti-HA Ab from the basolateral compartment after 12h (Figure 3D).

Given that the biosynthetic secretion supplies ~35% of apically transcytosed CFTR, apical-to-basolateral transcytosis accounts for ~65% of basolaterally delivered channels in CFBE (Figure 3B). To visualize CFTR apical-to-basolateral transcytosis, the channel was apically labeled by anti-HA Ab capture (3h, 37°C), followed by immuno-EM. Apically endocytosed CFTR-Ab complex was detectable at the basolateral PM (Figure 3E-F).

If CFTR is basolaterally missorted from apical endosomes, the round-trip transcytosis of apically labeled CFTR should be observed. To this end, we simultaneously exposed CFTR expressing CFBE to apical anti-HA Ab and basolateral HRP-conjugated secondary Fab at 37°C. We monitored the time-dependent appearance of the HRP activity at the apical PM (Figure 3G). The results confirmed that apical CFTR-anti-HA Ab complexes successively underwent apical-to-basolateral targeting, association with HRP-Fab, and then basolateral-to-apical transcytosis.

Jointly, these results suggest that constitutive apical transcytosis preserves CFTR apical polarity, which would be compromised in the absence of trancytosis. To test this conjecture, we inhibited cargo delivery to and retrieval from the basolateral PM by tannic acid (TA) crosslinking (Polishchuk et al., 2004). Basolateral TA exposure (15min, 37°C) accelerated the loss of apical CFTR to 11.3 ±1.2%/h compared to control (~5.5%/h) in dox-OFF CFBE (Figure 3H). This translates a reduction of CFTR half-life from ~12.4h to ~5.8h, assuming an exponential decay kinetic, in the absence of apical transcytosis and underscores the significance of transcytosis in CFTR apical PM stability.

### CFTR transcytotic route in airway epithelia

To delineate the transcytotic route of CFTR in CFBE, the channel was basolaterally labeled with anti-HA Ab (45min, 37°C), while HA detection at the cell surface was prevented with anti-mouse Fab. CFTR was then colocalized with organelle markers of ASE (EEA1), ARE (Rab11), CRE (Rab8), CRE and BSE (Transferrin, Tf), and fast recycling endosomes (Rab4) by indirect immunostaining. Transcytosed CFTR progressively accumulated in EEA1^+^ ASE and negligibly with Rab11^+^ ARE en route to the apical PM but failed to colocalize with Rab4 (Figures 4A, B and S3B). While no CFTR was colocalized with the CRE marker Rab8, we observed a modest but significant colocalization (~32%) with Cy3-Tf, especially in basal and middle sections of epithelia, while only ~19% of the TfR pool was colocalized with transcytosed CFTR throughout the cell (Figures 4A-B and S3C-D).

**Figure 4.**
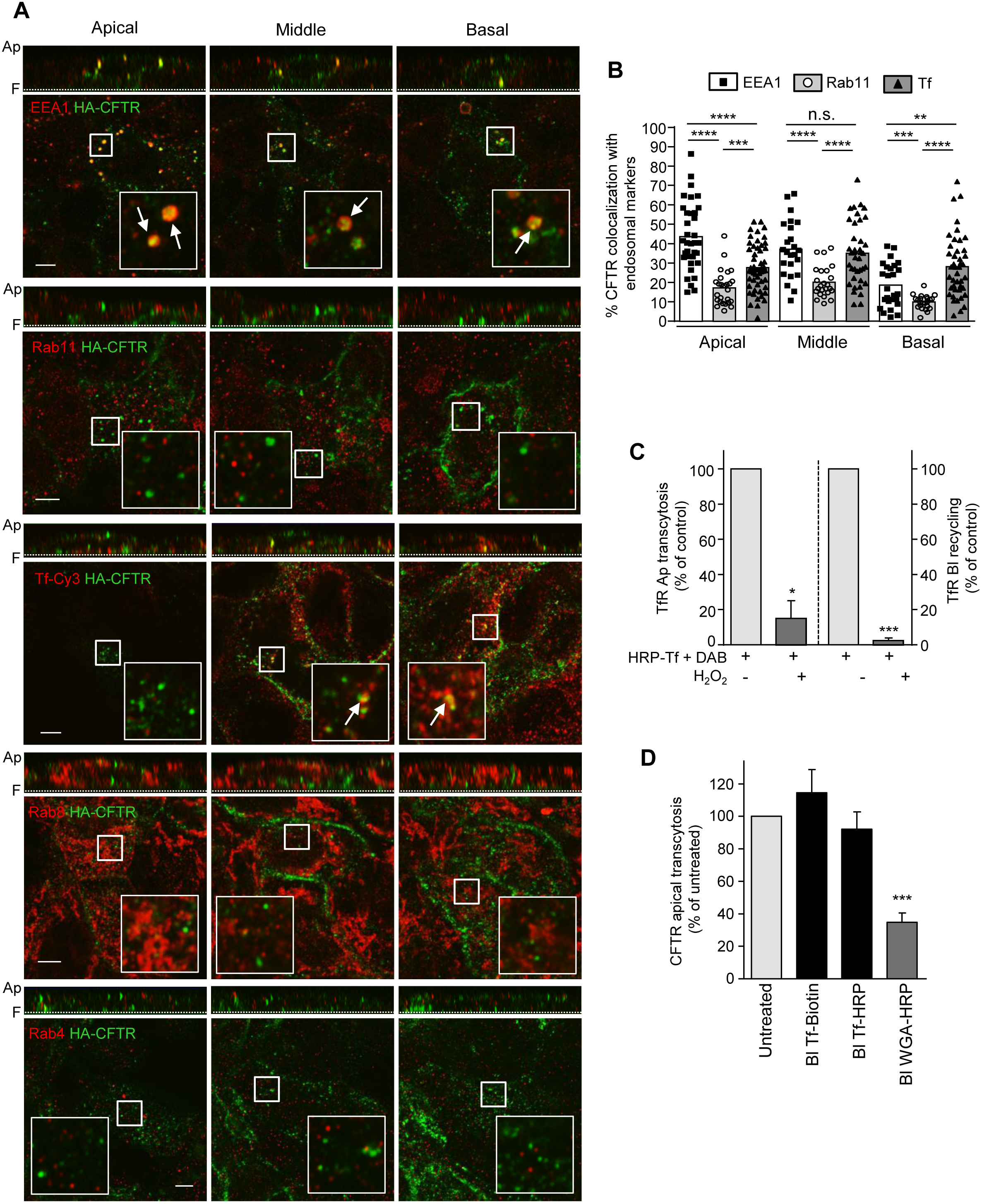
Apical transcytotic route of CFTR in polarized CFBE. (A) Immunocolocalization of transcytotic CFTR. CFTR was labeled by basolateral anti-HA Ab capture (45 min, 37°C), while staining of apically delivered CFTR-anti-HA complexes were blocked with goat anti-mouse Fab. Then, transcytotic CFTR-anti-HA (green) was colocalized with EEA1, Rab11, Rab4 or Rab8 (red) by indirect immunostaining and LCFM at the apical, middle, and basal planes of CFBE (lower panels). Vertical optical sections are shown as top panels. TfR was labeled by Cy3-Tf uptake (45min). Inserts represent ~3.5-fold magnification of the selected area. Ap, apical PM; F, filter. (B) Quantitative colocalization of transcytosed CFTR with organellar markers. Each dot represents the Mander’s coefficient of 23-54 R.O.I.s at the apical, middle or basal planes of CFBE (n=3, two-tailed unpaired t-test). (C) Functional ablation of TfR containing endosomes. Tf-HRP was basolaterally loaded (30 min, 37°C) and incubated with H_2_O_2_ and DAB or DAB alone (control). Apical transcytosis or basolateral recycling of Tf-HRP was measured as described in Methods and expressed as percentage of control (n=3). (D) CFTR apical transcytosis is largely independent of the TfR trafficking route but intersects with BSE. Following functional ablation of Tf-HRP or HRP-WGA loaded endosomes, CFTR apical transcytosis was measured for 3h at 37°C. Tf-biotin was used as control (n=4-5). Data are means ± SEM on each panel, * p<0.05, **p<0.01, *** p<0.001, **** p<0.0001.

To verify functionally whether transcytotic CFTR traverses TfR-containing endocytic compartments, basolateral endosomes were loaded with HRP-Tf (30min) to target the CRE (and to a much lesser extent the BSE) and these compartments were ablated using HRP-mediated cross-linking (Cresawn et al., 2007; Henry and Sheff, 2008). This protocol efficiently abrogated basolateral recycling and apical transcytosis of TfR but failed to alter CFTR transcytosis (Figure 4C-D). In contrast, ablation of BSE with HRP-WGA profoundly decreased CFTR transcytosis (Figure 4D). Overall, these results indicate that basolaterally internalized CFTR accumulates in EEA1^+^ ASE but seems to segregate from TfR^+^ CRE and Rab11^+^ ARE for apical transcytosis.

### Missorting of internalized CFTR for basolateral transcytosis

To confirm CFTR transcytotic itinerary in CFBE cells and assess the origin of CFTR loaded apical-to-basolateral transcytotic vesicles, we knocked down putative or proven regulators of CFTR endocytic traffic by siRNA (Figure S4A-B). Although ablation of Rab5 and Rab11 affected CFTR recycling (Gentzsch et al., 2004) (Figure 5A, top), they failed to influence CFTR transcytosis (Figure 5B). Similar results were observed after ablation of SNX4, a Rab11 interacting protein, ACAP1, a coat protein associated with recycling endosomes, as well as SNX17 which have been implicated in E-cadherin, Glut4 and integrin but not in CFTR recycling (Li et al., 2007; Solis et al., 2013; Steinberg et al., 2012) (Figure 5A-B). CFTR transcytosis remained also unaltered upon Rab4 and Rab8 depletion, confirming our immunostaining results (Figures 4A and 5B). Rab5 and ACAP1 siRNA ablation significantly, while SNX4 siRNA marginally decreased CFTR apical density, consistent with their permissive role in CFTR recycling (Figure 5A). These results suggest that basolateral missorting of internalized CFTR occurs before its Rab5-dependent entry into the ASE and confirm that CFTR apical transcytosis is independent of Rab8^+^ CRE and Rab11^+^ ARE in CFBE, in contrast to other transcytosed cargoes (e.g. pIgR), which transit the CRE and ARE prior to exocytosis (Jerdeva et al., 2010).

**Figure 5.**
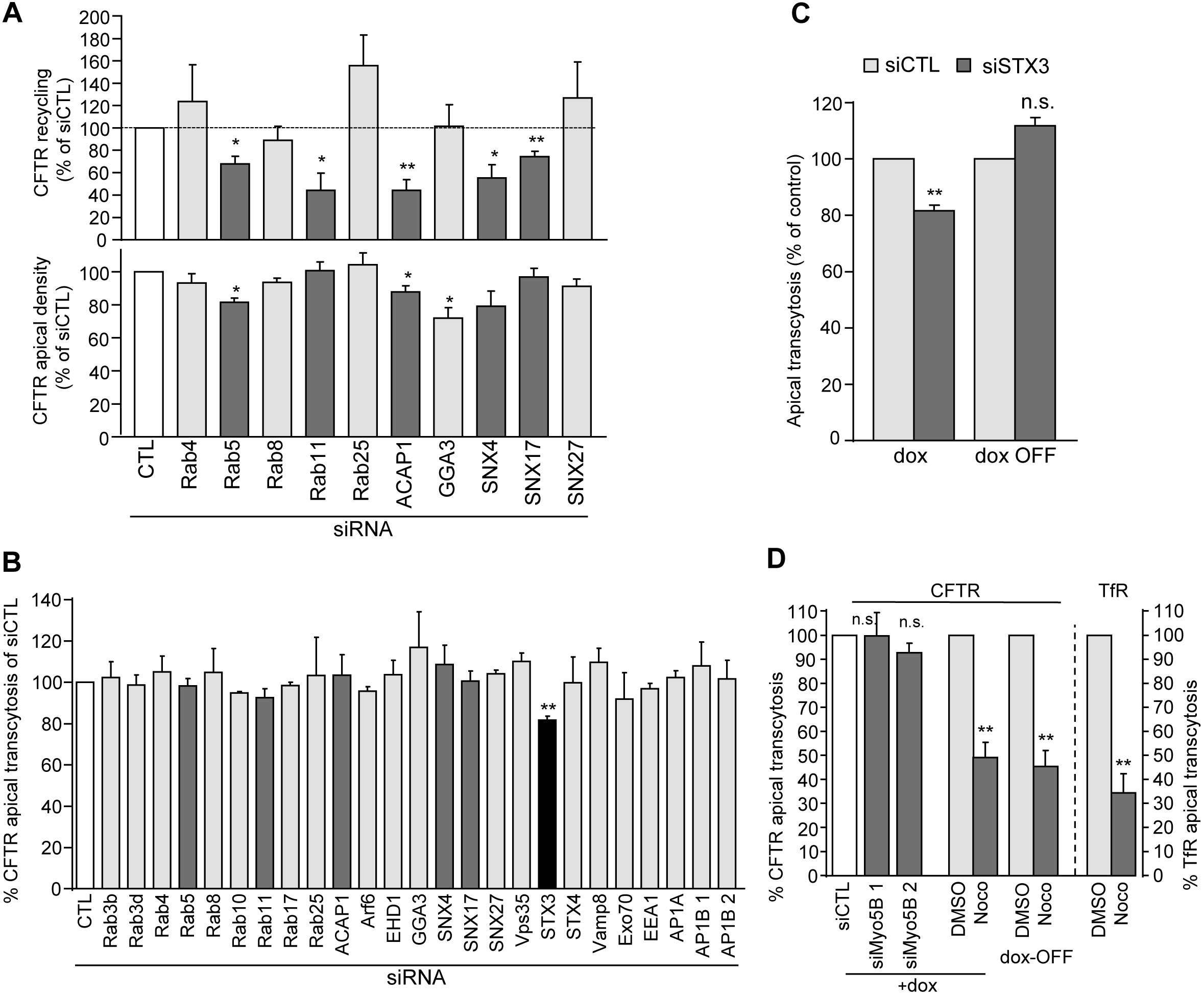
Modulation of CFTR transcytosis. (A-B) CFTR recycling, apical density and transcytosis were measured in CFBE silenced for the indicated protein. Results are expressed as % of the control siRNA (CTL) (n=2-42). (C) STX3 may contribute to the basolateral missorting of newly synthesized CFTR. CFTR apical transcytosis was measured after 24h dox-OFF in CFBE transfected with control (CTL) or STX3 siRNAs (n=3-4). (D) CFTR apical transcytosis is microtubules-dependent. CFTR and TfR transcytosis was determined in CFBE, transfected with control (CTL) or Myo5B specific siRNAs or pretreated for 30 min and during transcytosis with 33μM nocodazole or DMSO (n=3-6). Data are means ± SEM on each panel, n.s. non-significant, * p<0.05, ** p<0.01.

Only Syntaxin 3 (STX3) knockdown decreased CFTR apical transcytosis significantly (Figure 5B and S4A-B). Depletion or mutation of the apical targeting signal of this t-SNARE enhances basolateral mistargeting of NHE3 and p75-GFP (Sharma et al., 2006; Vogel et al., 2015). Interestingly, a fraction of STX3 was localized at the basolateral PM and its retrieval through an ubiquitin-dependent mechanism facilitated the recruitment of cargoes into apical exosomes (Giovannone et al., 2017). Considering that biosynthesis arrest abrogated the effect of STX3 siRNA on CFTR transcytosis (Figure 5C, dox-OFF), we suggest that STX3, besides controlling CFTR apical exocytosis (Collaco et al., 2010) may be involved in the delivery and/or retrieval of basolaterally targeted channels from the biosynthetic pathway.

### CFTR transcytosis requires microtubule

To test the contribution of myosinVB (Myo5B) and microtubules (MT), major carriers of transcytotic cargoes (Jerdeva et al., 2010; Tzaban et al., 2009), MTs were disrupted with nocodazole and Myo5B expression was silenced by siRNA. While Myo5B knockdown modestly decreased CFTR apical expression as a consequence of reduced entry of apically endocytosed channel into ARE (Swiatecka-Urban et al., 2007), it failed to interfere with apical transcytosis (Figures 5D and S4A,C). Nocodazole, however, profoundly decreased CFTR transcytosis similarly to TfR (Figures 5D and S4D). This cannot be attributed to the impeded biosynthetic transport of CFTR, as comparable inhibition of transcytosis (54.7 ±6.7% vs 50.9 ±6.3%) was observed in dox-OFF cells (Figures 5D and S4E). Considering that the channel endocytosis and recycling rates remained unaltered upon MT disruption (Figure S4F), the decreased CFTR apical expression could be attributed to its impeded transcytosis (13.4 ±4% in 3h) (Figures S4G and 5D).

### Transcytosis counteracts CFTR missorting upon disruption of PDZ protein(s) interaction

N- and O-glycans, GPI-anchors, and cytosolic and transmembrane segments have been identified as apical sorting determinants (Cholon et al., 2009; Kundu et al., 1996; Lisanti et al., 1989; Potter et al., 2006; Sharma et al., 2006). Basolateral targeting signals may partly overlap with internalization motifs of transmembrane cargoes, represented by tyrosine-(YXXΦ, NPXY) and di-leucine-based (D/EXXXLL) motifs (Stoops and Caplan, 2014).

To identify targeting motifs that may influence CFTR transcytosis efficacy, we inactivated three potential polarized sorting signals (Figure 6A). i) Three di-leucine and tyrosine-based endocytic motifs (Hu et al., 2001) were changed to alanine (K3-CFTR). ii) Two N-linked glycosylation sites (N894D- and N900D-CFTR) were inactivated in the 4th extracellular loop, iii) Considering the role of PDZ proteins in apical targeting, tethering and recycling of CFTR, we truncated the PDZ binding motif (DTRL) by eliminating the last six residues (Δ6-CFTR) at the C-terminus. All of these CFTR variants were stably expressed in CFBE (Figure S5A).

**Figure 6.**
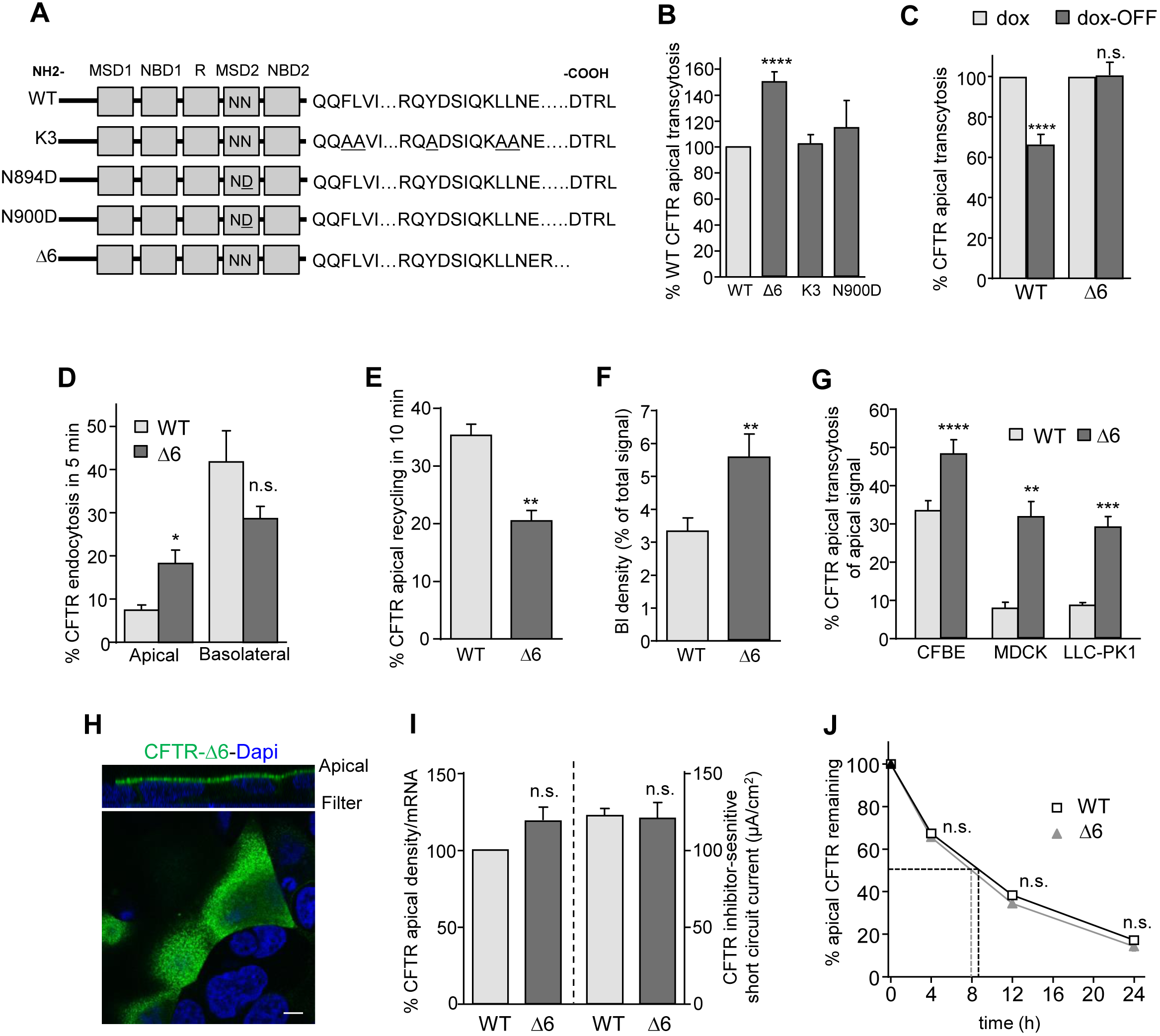
The effect of sorting signal and PDZ-domain binding motif mutation on CFTR polarized sorting and function in epithelia. (A) Mutagenesis of sorting signals that may influence CFTR transcytosis. Endocytic consensus sites ^1424^Tyr (Y) and ^1430-1431^Leu (L) were mutated to Ala (A). NN represents ^895^Asn and ^900^Asn two N-linked glycosylation sites in the MSD2. DTRL: PDZ-domain binding motif at the C-terminus. MSD, membrane spanning domain; NBD, nucleotide binding domain; R, regulatory domain. (B) Deletion of the DTRL motif (Δ6) increases CFTR apical transcytosis relative to WT-CFTR (n=3-16). (C) Biosynthetic basolateral missorting of CFTR depends on the binding of PDZ-domain protein(s). CFTR apical transcytosis was measured in control (+dox) or dox-OFF (after 24h) cells (n=17-18). (D-E) Apical but not basolateral CFTR internalization is accelerated upon deletion of the PDZ binding motif (n=3-7, two-tailed unpaired t-test). (E) Δ6 mutation impairs CFTR apical recycling (n=6), measured by PM-ELISA. (F) WT- and Δ6-CFTR basolateral cell-surface density was measured by PM-ELISA (n=11-15, two-tailed unpaired t-test). (G) The effect of Δ6 mutation on CFTR apical transcytosis in the indicated cell line (n=4-15). (H) Δ6-CFTR is predominantly detected at the apical PM in CFBE by indirect immunostaining, performed as in Figure 1D. Scale bar: 5μm. (I) The apical PM density and channel function of WT- and Δ6-CFTR were measured by PM-ELISA and 10μM forskolin-stimulated short circuit current (Isc), respectively, and corrected for their relative mRNA level (n=3). CFTR mediated Isc was quantified after inhibition with CFTR_inh_ 172. (J) Apical stability of WT- and Δ6-CFTR is similar. CFTR was labeled with anti-HA (1h, ice) and chased for the indicated time before PM-ELISA (n=4-6, two-tailed unpaired t-test). Data are mean ± SEM on each panel, n.s. not significant. * p<0.05, ** p<0.01, ***p<0.001, **** p<0.0001.

Surprisingly, apical transcytosis of Δ6-CFTR, but not K3- or N900D-CFTR, was increased by 49.5 ±7.8%, suggesting that the Δ6 mutation compromised the fidelity of CFTR apical biosynthetic targeting, endocytic recycling or both processes (Figure 6B). The comparable expression level of Δ6- and WT-CFTR-3HA and the inability of Δ6-, but not WT-CFTR to associate with the GST-NHERF1 fusion protein were confirmed (Figure S5B-C). Since transcytotic flux of Δ6-CFTR, but not WT-CFTR was insensitive to translation inhibition (Figure 6C, dox-OFF), the amplified basolateral arrival of Δ6-CFTR likely emanates from missorting at apical endosomes and not from the biosynthetic secretion. This inference is supported by the increased apical internalization (18.3 ±3.1 vs 7.4 ±1.2%/5min) and impeded recycling (20.5 ±1.8 vs 35.3 ±2.0%/10min) of Δ6-CFTR relative to WT-CFTR (Figures 6D-E). These data are also in line with CFTR tethering to the subapical actin cytoskeleton and enhanced channel recycling by PDZ domain proteins NHERFs (Haggie et al., 2004; Swiatecka-Urban et al., 2002), as well as with the ~1.5-fold increased basolateral density of Δ6-CFTR (Figure 6F). The accelerated basolateral transcytosis of Δ6-CFTR mirrors the stimulated apical transcytosis of TfR upon inhibition of its basolateral recycling by AP-1B ablation (Perez Bay et al., 2013). The apical transcytosis of Δ6-CFTR was also augmented in MDCK and LLC-PK1 (Figure 6G), supporting the notion that PDZ protein interaction(s) enhance the fidelity of CFTR polarized sorting/recycling in various epithelia.

Remarkably, Δ6-CFTR displayed similar biochemical and functional apical PM expression as WT-CFTR despite its accelerated internalization and impeded apical recycling, measured by immunostaining, PM ELISA and short circuit current (Isc), respectively (Figure 6H, I). Finally, we could not resolve a detectable difference in the PM turnover of Δ6- and WT-CFTR by PM-ELISA or biotinylation assay (Figure 6J and S5D), suggesting that 2-fold augmented round-trip transcytosis maintains the WT-like apical PM stability of Δ6-CFTR and highlights the sorting capacity and significance of transcytotic pathway to preserve CFTR polarity.

### NHERF1 association attenuates CFTR basolateral missorting and transcytosis

To search for additional proteins that may contribute to the recycling fidelity of apically internalized CFTR, the consequence of siRNA ablation of CFTR-interacting PDZ domain proteins was measured (Figures S4B and S5F). The apical transcytosis of WT- but not Δ6-CFTR or TfR was increased only by NHERF1 (EBP50) knockdown (Figure 7A-B). NHERF1 siRNA accelerated the internalization and inhibited the recycling of WT-, but not Δ6-CFTR, at moderately altered CFTR apical density (Figure 7C-E). Finally, we showed that NHERF1 was partly colocalized with internalized CFTR in a subapical compartment (Figures 7F-G and S5G-H). These complementary results reinforce the functional relevance of NHERF1-CFTR interaction in the channel apical retention and endosomal recycling along the ASE/ARE compartments in CFBE and provide a plausible explanation for the exacerbated basolateral missorting of CFTR upon preventing NHERF1-CFTR association either by C-terminal truncation of CFTR or NHERF1 ablation.

**Figure 7.**
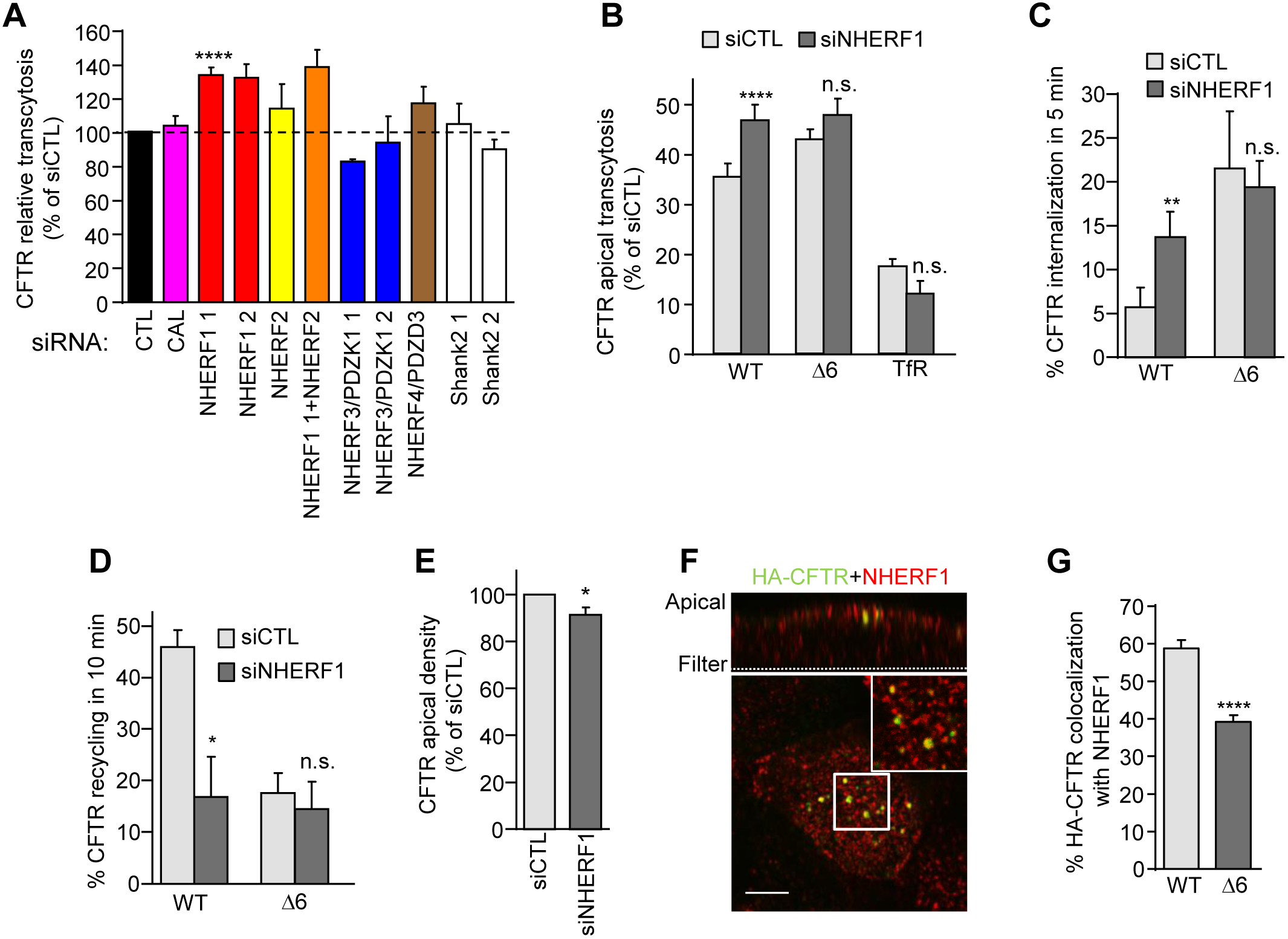
NHERF1 ablation phenocopies the Δ6-CFTR cellular phenotype. (A) CFTR transcytosis was assayed after siRNA-mediated knockdown of the indicated PDZ proteins in CFBE and expressed as the % of siCTL treated cells (n=2-19). (B-E) NHERF1 silencing mirrors the PDZ-binding motif truncation of CFTR. CFBE were transfected with control siCTL or NHERF1 1 siRNA and then assayed for (B) CFTR and TfR transcytosis following 3h basolateral anti-HA or Tf-HRP capture (n=3-10), (C) CFTR internalization (n=5-6), (D) CFTR recycling (n=3-4), (E) CFTR apical PM density (n=10), measured as described in Methods. (F) NHERF1 colocalizes with the apical endocytosed CFTR. CFTR was labeled by anti-HA capture (1h, 37°C) from the apical compartment. Residual CFTR apical PM was blocked with goat anti-mouse Fab on ice. NHERF1 (red) and CFTR (green) were visualized by indirect immunostaining and LCFM in permeabilized cells. Upper panel represents a z-section. The insert is a 4-fold magnification of the R.O.I. Scale bar: 5 μm. (G) Quantification of colocalization between NHERF1 and subapical endocytosed WT- and Δ6-CFTR in CFBE cells (n=42-52 R.O.I., 2 independent experiments, two-tailed unpaired t-test). Data are mean ± SEM on each panel, * p <0.05, ** p<0.01, **** p<0.0001.

## DISCUSSION

Here we report that apical and basolateral transcytosis represent previously unrecognized membrane trafficking pathways that significantly contribute to the development and maintenance of CFTR apical polarity, as well as the phenotypic suppression of CFTR C-terminal truncations in multiple epithelia, including immortalized and primary human airway. We show that both CFTR apical transcytosis and highly efficient basolateral internalization play a role in counteracting the channel constitutive basolateral missorting and maintaining ~30-fold higher amount of CFTR at the apical than at the basolateral PM in CFBE (Figures S5E and S6). The reduced rate of CFTR transcytosis in MDCK cells and the limited efficiency of CFTR biotinylation relative to ELISA-based assay may explain that CFTR (or EGFR) transcytosis was undetectable previously (Cotton et al., 2013; Swiatecka-Urban et al., 2002).

The physiological significance of CFTR apical transcytosis is illustrated by the >2.1-fold reduction of its half-life at preserved apical internalization upon inhibition of basolateral retrieval. CFTR basolateral accumulation in the absence of apical transcytosis would decrease apical Cl^-^ secretion by reducing basolateral Cl^-^ entry, causing attenuated coupled water secretion and mucociliary clearance of airway epithelia (Ballard et al., 2002; Farmen et al., 2005).

We also demonstrate that CFTR basolateral delivery predominantly (~65%) originates from apical endocytic vesicles via reversed transcytosis, due to limited fidelity of its apical recycling. A similar phenomenon prevails for endosomal missorting of TfR and EGFR at basolateral endosomes (Cotton et al., 2013; Gravotta et al., 2007). In addition, missorting of newly synthesized CFTR from the TGN fuels ~35% of the channel basolateral delivery, likely in a STX3-dependent manner.

Based on colocalization and functional endosomal ablation studies, we propose that basolaterally internalized CFTR enters transcytotic endosomes (TE), which are EEA1^+^ with reduced TfR content, that fuel the transcellular migration of CFTR in a MT-dependent, but Myo5B-independent pathway (Figure S6). Several observations suggest that CFTR apical transcytosis has some unique characteristics in CFBE: i) Unlike basolateral sorting of TfR, which is thought to proceed at CRE (Gravotta et al., 2012), commitment of CFTR to apical recycling likely takes place before its Rab5-mediated entry into ASE. This inference is supported by the observation that accelerated endocytosis of CFTR lacking its PDZ binding motif diplays enhanced basolateral targeting and subsequent apical transcytosis, whereas silencing of proteins along CFTR endocytic sorting (Rab5, EEA1 or SNX17) or recycling (Rab11, ACAP1, or SNX4) failed to do so. ii) The results of CFTR mutagenesis and siRNA screens showed that transcytosis of CFTR in CFBE is independent of N-glycans, AP-1B, Rab3b, Rab17, Rab25 or the exocyst complex that are essential for the apical transcytosis of pIgR, TfR and FcRn (Hunziker and Peters, 1998; Nelms et al., 2017; Perez Bay et al., 2014; Perez Bay et al., 2013; Tzaban et al., 2009; van et al., 2002). iii) Finally, transcytotic CFTR seems to avoid the CRE and ARE compartments that are commonly used by other transcytotic cargoes (Jerdeva et al., 2010; Lalioti et al., 2016). Interestingly, constitutive TfR apical transcytosis in AP-1B-expressing MDCK (Perez Bay et al., 2013) or CFBE (Figures S3C-E and S4H) also avoids the Rab11^+^ ARE, similar to that of FcRn (Tzaban et al., 2009). Hence, it is plausible that in some epithelia the transcytotic pathway of CFTR and other cargoes intermix in a common endosomal compartment prior to apical exocytosis that is distinct from the ARE. Indeed, sorting endosomes are composed of unique microdomains that sort cargoes with different mechanisms (i.e. PDZ motif- and actin-dependent recycling of β_2_ adrenergic receptor (β_2_-AR) and bulk recycling of TfR (Puthenveedu et al., 2010)). This would explain the modest colocalization between CFTR and Tf at subapical pole of CFBE, as well as the EEA1- and Rab-5-independent CFTR transcytosis, despite the channel residence in EEA1^+^ endosomes.

Comparison of the exocytotic machinery of apical CFTR with other transcytotic cargoes derived from the biosynthetic, recycling, and transcytotic pathways of airway epithelia remains to be investigated. Intriguingly, we failed to detect a significant reduction of CFTR apical transcytosis and PM density upon ablation of Myo5B, which has been implicated in maintaining CFTR apical expression along recycling and secretory pathways in CFBE and CaCo2 cells (Swiatecka-Urban et al., 2007; Vogel et al., 2015). Recent studies also failed to detect profound changes in CFTR functional expression in intestinal epithelia expressing either an inactive Myo5B or following Myo5B knockdown (Kravtsov et al., 2016). Furthermore, Myo5B-independent apical polarization of CFTR is in line with the augmented Cl^-^ secretion in Microvillus inclusion disease, caused by loss-of-function of Myo5B (Kravtsov et al., 2016).

Our results expand the established role of NHERF1 in CFTR traffic at multiple cellular locations. i) By tethering CFTR to the subapical cytoskeleton, NHERF1 enhances the channel apical retention (Haggie et al., 2004; Swiatecka-Urban et al., 2002). Importantly, this mechanism is absent at the basolateral PM, due to considerably reduced concentration of NHERFs, ezrin and cortical actin (Reczek et al., 1997; Short et al., 1998), accounting for CFTR fast internalization and limited recycling, ensuring its low basolateral PM density. ii) We provide evidence for enhanced colocalization of NHERF1 with the endocytic pool of WT-CFTR relative to that of Δ6-CFTR (Figure 7F, G), supporting the paradigm that NHERF1-mediated sorting directs the channel towards recycling at subapical endosomes (ASE and/or ARE) (Cardone et al., 2015; Cushing et al., 2008). iii) Finally, we demonstrate that PDZ motif elimination has negligible influence on CFTR apical expression, transport activity and stability. Increased basolateral internalization and apical transcytosis flux of basolateral CFTR counteract the channel accelerated removal and decreased recycling at the apical PM. These results expand previous observations obtained on CFTR variants with compromised binding of PDZ proteins (Benharouga et al., 2003; Ostedgaard et al., 2003) and explain the isolated elevation of sweat Cl^-^ concentration in the absence of pancreatic and lung phenotype in individuals harboring a single copy of C-terminal 26 residues of CFTR (S1455X-CFTR) (Mickle et al., 1998). Finally, upregulated CFTR compensatory transcytosis may also explain the mild CF cellular phenotype (Ameen et al., 2007) of the naturally occurring N287Y-CFTR mutant that causes accelerated apical internalization (Silvis et al., 2003).

Whether CFTR transcytotic routes are affected by CF-causing mutations and can influence a non-native channel degradation via ubiquitin-dependent lysosomal targeting from the PM, such as ΔF508 (Okiyoneda et al., 2018), remains to be explored. Interestingly, a CF-causing folding mutant, P67L-CFTR (Bagdany et al., 2017; Sabusap et al., 2016), displays considerably reduced transcytotic flux that can be restored to that of WT upon folding correction with VX-809 (data not shown). Intriguingly, overexpression of NHERF1 or ablation of CAL (CFTR <u>a</u>ssociated ligand) that facilitates CFTR lysosomal targeting at the TGN partly rescues the PM expression of ΔF508 (Guerra et al., 2005; Wolde et al., 2007). Considering that deletion of the PDZ motif decreased CFTR biosynthetic delivery to the basolateral PM (Fig. 6C, dox-OFF), we speculate that NHERF1 overexpression may rescue ΔF508 not only by increasing its retention/recycling at the apical PM but also by promoting its biosynthetic over endocytic basolateral missorting, which may further reduce the degradation propensity of the channel by reducing basolateral transcytosis.

## MATERIALS AND METHODS

### Antibodies and reagents

Antibodies used in this study are listed in Supplementary Table 1. Nocodazole, MESNA (Sodium 2-mercaptoethanesulfonate), DAB (3,3’-diaminobenzidine tetrahydrochloride), Horseradish peroxidase (HRP) and tannic acid were purchased from Sigma-Aldrich. Y-27632 was from Tocris. HRP-Tf and Ultroser-G were purchased at Jackson Immunoresearch and Pall Corporation, respectively.

### Cell culture and medium

CFBE, Calu-3 and 3T3-J2 cells are generous gifts from D. Gruenert (University of California-San Francisco, USA), J.W. Hanrahan (McGill University, Canada) and B. Scholte (Erasmus MC, Rotterdam, Netherlands), respectively. LLC-PK1 and NCI-H441 were purchased from the ATCC. MDCKII was described before (Benharouga et al., 2003; Veit et al., 2012). CFBE cells were propagated in MEM medium supplemented with fetal bovine serum (FBS), L-Glutamine and HEPES (Invitrogen) on coated plastic flasks as described before (Veit et al., 2012) and were seeded and differentiated for ≥3 days on coated plastic wells or polyester permeable supports (Transwell filters, Corning). MDCK II and LLC-PK1 and NCI-H441 epithelial cells were cultured in DMEM (Invitrogen) supplemented with 10% FBS. Human lung adenocarcinoma Calu-3 cells were cultured in DMEM/F12 (Invitrogen) supplemented with 10% FBS. All cells were maintained in a 37°C incubator under 5% CO_2_. CFBE, MDCK, NCI-H441, and LLC-PK1 cell lines, expressing inducible WT- and Δ6-CFTR with a 3HA or HRP tag were generated using the ClonTech pLVX-Tight-Puro lentivirus technology, as described previously (Veit et al., 2014; Veit et al., 2012; Veit et al., 2018) and induced with 250-500ng/ml dox. MDCKII cells, stably expressing WT- or Δ6-CFTR-3HA, were generated by transduction with retroviral particles (Benharouga et al., 2003).

### CR-HBE characterization and expression of CFTR-3HA

Primary human bronchial epithelial cells (HBE) cells were isolated in W.E. Finkbeiner’s laboratory under approval from the University of California, San Francisco Committee on Human Research as well as purchased from the Cystic Fibrosis Translational Research center (CFTRc), McGill University. HBE were conditionally reprogrammed and differentiated according to a modified protocol of Liu et al. (Liu et al., 2012). Briefly, HBE cells were cultured on irradiated 3T3-J2 fibroblasts in proliferation F-Medium (Liu et al., 2012) with 10μM of ROCK inhibitor Y-27632. After expansion, cells were plated on collagen IV coated transwell filters and differentiated in Ultroser-G medium (Neuberger et al., 2011) at air-liquid interface during at least 4 weeks with basolateral medium change every 2-3 days. CR-HBE cells constitutively expressing WT-CFTR-3HA were developed by transducing the cells with lentiviral particles encoding the WT-CFTR-3HA in pTZV4-CMV-IRES-puro (Open Biosystems) as described (Veit et al., 2012) during proliferation, followed by two days puromycin selection before seeding on filter supports. Air Surface Liquid (ASL) height was determined after staining the ASL with 2 mg/ml tetramethyl-rhodamine dextran (D18-19, 70,000 MW, Neutral, Invitrogen), dispersed in a low boiling point perfluorocarbon (Fluorinert FC-72, boiling point 56°C, 3M Company) and the epithelia with Cell tracker (Invitrogen). The ASL height was determined in 3-10 areas of 2 filters using a Zeiss700 upright fluorescence laser confocal microscope equipped with an environmental chamber at 37°C and 5% CO_2_.

### CFTR cell surface ELISA and transcytosis assay

CFTR apical density and apical transcytosis were measured by PM-ELISA. Cells were incubated with anti-HA (1:1000) in the apical (apical density) or basolateral (transcytosis) compartment, respectively, at 37°C for 0.5-4 h. After washing with ice-cold PBS (Gibco) supplemented with 1 mM MgCl_2_ and 0.1 mM CaCl_2_ (PBSCM) of the apical PM, all the cells were incubated apically with horseradish peroxidase (HRP)-conjugated anti-mouse Ab (1:1000) in PBSCM-0.5% bovine serum albumin (PBSCM-BSA) (1h, on ice). Cells were washed with PBSCM and the HRP activity in the apical compartment was determined in the presence of Amplex Red fluorogenic substrate (Invitrogen) at 544 nm excitation and 590 nm emission wavelengths, using a POLARstar OPTIMA (BMG Labtech) or a Tecan Infinite M1000 (Tecan Group) fluorescence plate reader. CFTR apical transcytosis was expressed relative to its apical density, measured in parallel (% Ap transcytosis = (Ap signal^transcytosis^/Ap signal^ap density^)*100). Non-specific Ab binding was determined by using either non-expressing CFBE cells or replacing the anti-HA with non-specific mouse IgG under the same experimental conditions. EGFR transcytosis was measured using the same PM ELISA assay, with anti-EGFR instead of anti-HA. To monitor TfR apical transcytosis, endogenous Tf was first depleted in serum-free medium (1h), before incubation of HRP-Tf or HRP (both 5ug/ml) in the apical or basolateral compartment. After 3h, the peroxidase activity was measured on the apical PM and in the medium. TfR transcytosis was expressed as the signal intensity in the apical compartment relative to that of TfR steady state density at the basolateral PM. Round-trip transcytosis was measured by incubating simultaneously the cells with mouse anti-HA (1:1000) in the apical chamber and with HRP-conjugated anti-mouse Fab (1:1000) (Jackson Immunoresearch) in the basolateral chamber for 4 to 24h at 37°C. Cells were then extensively washed with ice-cold PBSCM and assessed for HRP activity in the apical compartment.

### CFTR internalization and recycling assays

CFTR internalization rate was measured by determining the CFTR-Ab complexes disappearance from the apical PM by ELISA, as described earlier (Glozman et al., 2009). Apical CFTR was labeled with anti-HA (1:1000) for 1h on ice. After washing with ice-cold PBSCM, one plate was incubated for 5 min at 37°C (P5’), while the control plate was kept on ice (P0). The amount of CFTR remaining at the PM was determined using HRP-conjugated secondary Ab and Amplex Red in both plates. The internalization rate was expressed as the percentage of the initial amount of CFTR removed in 5 min (% CFTR internalization = (P0-P5’)*100/P0).

CFTR recycling rate was assessed by monitoring the exocytosis of internalized CFTR-Ab complexes at the apical PM. To this end, CFTR expressing CFBE cells were seeded on 4 plates (P1-P4). After anti-HA binding to apical CFTR (1h, on ice, P1-P4), the CFTR-Ab complex was internalized for 30 min at 37°C (P2-P3-P4) while a control plate was kept in cold medium (P1). The PM remaining anti-HA was blocked with anti-mouse monovalent Fab (Jackson ImmunoResearch, 1:75, 1h, on ice, P3-P4). Recycling of endocytosed CFTR was elicited at 37°C for 10 min (P4) while a blocking efficiency plate was kept in cold medium (P3). All plates were apically incubated with HRP conjugated anti-mouse Ab (1:1000) for 1h on ice, washed and peroxidase activity was measured. The recycling efficiency in 10 min was expressed as the percentage of endocytosed pool of CFTR, taking into consideration the blocking efficiency of the anti-mouse Fab (%Ap recycling = (P4-P3)*100/(P1-P2)).

### Time course of polarized delivery of newly synthesized CFTR-HRP

To measure newly translated CFTR delivery kinetics to the apical and basolateral PM, CFTR-HRP expression was induced for ~3.5h at 37°C. Then cells were washed with buffer H (154 mM NaCl, 3 mM KCl, 10 mM Hepes, 1mM MgCl_2_, 0.1 mM CaCl_2_, 10 mM glucose, pH 7.8) and incubated further in buffer H, complemented with Amplex Red at 37°C. Apical and basolateral media were sampled for Amplex Red fluorescence in every 5 min for ~20 min. Non-induced cells were used to determine the background fluorescence.

### Basolateral PM turnover inhibition by tannic acid

The basolateral membrane turnover was inhibited by tannic acid, using a modified protocol of Polishchuk et al. (Polishchuk et al., 2004). CFBE cells were treated for 15 min with 0.5% tannic acid in serum-free medium, then washed 2 times in serum-free medium and once in full medium followed by PM ELISA. Reversibility of the tannic acid treatment was ascertained by measurement of CFTR-anti-HA complex basolateral uptake after 8h chase (data not shown).

### Inactivation of endosomal compartments by HRP-mediated ablation

Tf-containing endosomes were ablated as described previously (Cresawn et al., 2007) with the following modifications. Tf-depleted CFBE cells were loaded with HRP-Tf (5 μg/ml) for 30 min (37°C) from the basolateral chamber. Cells were then washed with ice-cold PBSCM and incubated 2×5 min with 150 mM NaCl and 20 mM citric acid, pH 5.0 to remove cell surface-bound HRP-Tf. Wheat germ agglutinin (WGA)-labeled endosomes were inactivated after 20 min internalization of WGA-HRP (10 μg/ml, Sigma) from the basolateral compartment. WGA-HRP remaining at the cell surface was removed with 3×10 min incubation with 100 mM N-acetyl-D-glucosamine. Endosomal ablation was induced by the addition of 0.1 mg/ml DAB and 0.025% H_2_O_2_ for 1h on ice in the dark. The reaction was quenched with PBSCM-1% BSA. As controls, DAB or H_2_O_2_ were omitted, or Tf-biotin and WGA-Alexa 647 were used instead of Tf-HRP and WGA-HRP, respectively.

### siRNA-mediated inhibition of gene expression

Stealth siRNAs and siRNAs were purchased from Invitrogen, Origene and Qiagen, respectively. To limit off-target effects, the siRNAs were used at a final concentration of 25nM and phenotypic screens were carried out using pools of two to four siRNAs. For filter-based experiments, CFBE cells were electroporated using the Neon Transfection system (Invitrogen) with three 10 ms 1500V pulses using 100 μl Neon pipette tips according to manufacturer’s protocol. Cells on coated plastic were transfected with Lipofectamine RNAiMax (Invitrogen) according to the manufacturer’s protocol and then were polarized for 4 days in presence of dox-induced CFTR expression.

### Immunofluorescence microscopy and colocalization

Differentiated filter-grown CFBE cells were fixed in 4% paraformaldehyde and permeabilized with 0.2% Triton X-100. After blocking in PBSCM-BSA 0.5%, cells were incubated with primary Ab, washed with PBSCM and incubated with Alexa Fluor-conjugated secondary Abs. Extensively washed and cut-out pieces of filters were mounted between glass slide and coverslip. For monitoring intracellular CFTR colocalization, CFTR was labeled by anti-HA capture at 37°C for the indicated time. When indicated, PM CFTR staining was blocked with anti-mouse Fab prior to fixation (1h, on ice). TfR was visualized by exposing the cells to Cy3-Tf (20 μg/ml, Jackson Immunoresearch, 45min-1h at 37°C) in the basolateral compartment after depletion of the endogenous Tf in serum-free medium (1h, 37°C). Nuclei were stained with DAPI. 20-30 horizontal optical sections were acquired using a LSM-710 or LSM-780 LFCM equipped with a Plan-Apochromat 63X/1.40 oil objective (Carl Zeiss), reconstituted using the Zen 2012 software package and representative vertical sections are shown. For colocalization study, Mander’s coefficient was calculated using the JACOP plugin of ImageJ software after background subtraction and thresholding from individual optical sections at the apical, middle and basal plane.

### Monitoring CFTR apical stability by cell surface biotinylation and immunoblotting

Apical surface proteins of filter-grown CFBE cells were biotinylated for 15 min on ice with 1 mg/ml EZ Link sulfo-NHS-SS-biotin (Thermo Fisher Scientific) in buffer H (154 mM NaCl, 3 mM KCl, 10 mM Hepes, 1mM MgCl_2_, 0.1 mM CaCl_2_, 10 mM glucose, pH 7.8). Excess biotin reagent was quenched with 100 mM glycine in PBSCM. Cells were then shifted to 37°C for 0, 4, 12 or 24 h before being lysed in RIPA buffer (150 mM NaCl, 20 mM Tris-HCl, 1% Triton X-100, 0.1% SDS, and 0.5% sodium deoxycholate, pH 8.0) containing proteases inhibitor. Biotinylated proteins were isolated from postnuclear lysates with streptavidin-agarose beads (Invitrogen) at 4°C with end-over-end rotation. Proteins were eluted with 5X Laemmli sample buffer and visualized by immunoblotting with anti-HA or a mixture of L12B4 and M3A7 mouse monoclonal anti-CFTR Abs followed by either IRDye 800 anti-mouse Abs (Licor Biosciences, Lincoln, NE, USA) with the Odyssey Infrared Imaging System (Licor Biosciences) or using enhanced chemiluminescence as described (Benharouga et al., 2001).

### Electron microscopy

Filter-grown CFBE cells were incubated with anti-HA (1:500) in the basolateral or apical compartment for 3h at 37°C. Cells were washed, fixed, permeabilized and incubated with 1.4nm nanogold-conjugated anti-mouse Fab fragment (1:50) (Nanoprobes, Yaphank, NY, USA) for 1h on ice. Cell monolayers were then post-fixed with 1% osmium tetroxide in 0.1 M phosphate buffer for 1h at 4°C. Samples were dehydrated through a series of graded ethanol baths and embedded in epon resin. Semi-thin sections of ~1μm were obtained with a diamond knife on an ultramicrotome (Ultracut E, Reichert-Jung) and stained with 1% toluidine blue. Then, 60 nm thick sections were cut and counterstained with 4% uranyl acetate and Reynold’s lead citrate. Sections were observed under a Philips CM120 electron microscope equipped with a Gatan digital camera. Nanogold particles per μm of PM were calculated using the ImageJ software. Post-labeling steps and imaging were performed at the EM facility of the Department of Pharmacology, McGill University.

### Short circuit current (Isc) measurement

Isc of CFTR expressing CFBE was measured on cells differentiated on 12 mm Snapwell filters (Corning) mounted in Ussing chambers and bathed in Krebs-bicarbonate buffer (140 mM Na^+^, 120 mM Cl^-^, 5.2 mM K^+^, 25 mM HCO3^-^, 2.4 mM HPO_4_, 0.4 mM H_2_PO_4_, 1.2 mM Ca^2+^, 1.2 mM Mg^2+^ and 5 mM glucose, pH 7.4) at 37°C in the presence of an apical-to-basolateral chloride gradient. To functionally isolate apical PM, the contralateral domain was permeabilized with 100 μM amphotericin B. In each experiment, 100 μM amiloride and 20 μM forskolin were added sequentially to the apical and basolateral side to inhibit the epithelial Na^+^ channel and activate CFTR, respectively. CFTR activity was inhibited by 20 μM of channel blocker CFTR_inh_ 172.

### Recombinant NHERF1 purification and pulldown

GST or GST-NHERF1 was expressed in BL21 bacterial strain by pGEX-4T plasmid. Bacterial pellets were resuspended in 50 mM Tris-Cl pH 8, 50 mM NaCl, 5 mM EDTA, 0.5% NP-40, 5% glycerol (50 μl/ml) and sonicated. The lysate was centrifuged at 14000x rpm (30 min, 4°C) and passed through a Dowex 50X2-400 ion-exchange resin (Acros Organics). The flow through was incubated with glutathione-sepharose 4B beads (GE Healthcare) for 2 h at 4°C. After 3 washes, beads were incubated with the post-nuclear supernatant of the CFBE cell lysate, expressing WT- or Δ6-CFTR for 2h at 4°C under rotation. After 3 washes with RIPA, co-isolated CFTR was immunoblotted.

### Metabolic pulse labeling

Post-confluent filter-grown CFBE cells were incubated with methionine- and cysteine-free alpha-MEM medium for 45 min at 37°C. Cells were then pulse labeled in the presence of [^35^S]-methionine and [^35^S]-cysteine (0.1 mCi/ml; Perkin Elmer, Waltham, MA, USA) from the basolateral compartment for 30 min at 37°C in a humid chamber. After washing with ice-cold PBSCM, CFTR was immunoprecipitated with a mixture of M3A7 and L12B4 anti-CFTR Abs. Following autoradiography, radioactivity incorporated into CFTR was quantified by phosphorimage analysis, using a Typhoon workstation (GE Healthcare).

### EndoH and PNGase F digestion of CFTR

CFBE and Calu-3 cells expressing CFTR-3HA and endogenous CFTR were grown on 6 cm coated plastic dishes for 4-5 days post confluency. CFTR expression in CFBE was induced with 250 ng/mL dox for 4 days. Cells were lysed (0.3 % Triton X100, 150 mM NaCl, 20mM Tris pH 8.0) and after centrifugation (at 4°C, 12000xrpm for 10 min), the supernatants were digested with EndoH or PNGase F enzyme according to the manufacturer’s protocol. Samples were immunoblotted with anti-CFTR antibodies (L12B4, M3A7).

### RT-qPCR

For WT- and Δ6-CFTR mRNA expression, total RNA was extracted from CFBE lysed in Qiazol and analyzed using the one-step QuantiFast SYBR Green RT-PCR Kit (Qiagen, 204154) as recommended by the manufacturer. Briefly, total RNA was extracted from polarized CFBE grown on coated plastic in 24 well plates using the miRNeasy Mini Kit (Qiagen, 217004). Reverse transcription and PCR amplification was done sequentially in a Stratagene Mx3005P real-time thermocycler (Agilent, 401513) during the same thermocycler protocol on 50-100ng total RNA, determined by Nanodrop UV-Vis light absorbance. The abundance of transcripts was determined using a SYBR Green fluorescence amplification curve and its intersection with a preset threshold, yielding a Ct value. Data were analyzed by efficiency-corrected comparative quantification with MxPro QPCR software (Agilent) and the variations in initial RNA loading amount was normalized with GAPDH as a reference gene. mRNA expression differences between samples were reported as percent abundance relative to a reference sample (e.g. WT-CFTR). PDZ proteins down-regulation was evaluated with Quanti-Tect reverse transcription kit (Qiagen) as previously described (Veit et al., 2012).

### Statistical analysis

Results are presented as mean ± SEM of the number of independent experiments indicated in the figure legends, as biological replicates. Unless specified, P values were calculated with the means of at least 3 independent experiments by two-tailed paired student’s t-test and a 95% confidence level was considered significant. Normal distribution of data and homogeneity of variance were validated by calculating the skew factor (−2 > skew < 2) and performing the F-test, respectively. For non-normal data, Mann-Withney U-test was used for calculating the P-values and was indicated in the figure legend. For normal distributions with non-homogenous variances, the Welch correction was applied to two-tailed unpaired t-test to calculate the P-values and was indicated in the figure legend. All ELISA-based assays were performed using 2-4 technical replicates, except CFTR-HRP polarized delivery (1-2 technical replicates). Short circuit current measurement and qPCR were performed with 2 technical replicates.

## AUTHOR CONTRIBUTIONS

A.B.M., F.B. and G.L.L conceived the study. A.B.M., F.B., A.S., R.F., G.V. and H.X. performed and analysed the experiments. A.B.M. and G.L.L. wrote the manuscript with contributions from all authors.

## ACKNOWLEDGEMENTS

We are grateful to J.W. Hanrahan, B. Scholte and to the late D. Gruenert for cell lines and W. E. Finkbeiner for primary HBE. We thank R. Robert and R.G. Avramescu for help with intial Isc and qPCR measurements, respectively. Post-labeling steps and EM imaging were performed at the EM facility of the Department of Pharmacology, McGill University. The core facility of W.E. Finkbeiner (Department of Pathology, University of California, San Francisco) was supported by grants from National Institute of Health [DK072517] and Cystic Fibrosis Foundation Research and Translational Core Center [VERKMA15R0]. This work was supported by the Canadian Institutes of Health Research, NIH-NIDDK, and Cystic Fibrosis Canada to G.L.L. A.B.M. was supported by a travel grant from Cystic Fibrosis Canada. F.B. was a recipient of a Richard & Edith Strauss Fellowship, McGill Univeristy. G.L.L. is a Canada Research Chair.

The authors declare no competing financial interests.

## SUPPLEMENTARY INFORMATION

**Supplementary Figure 1.**
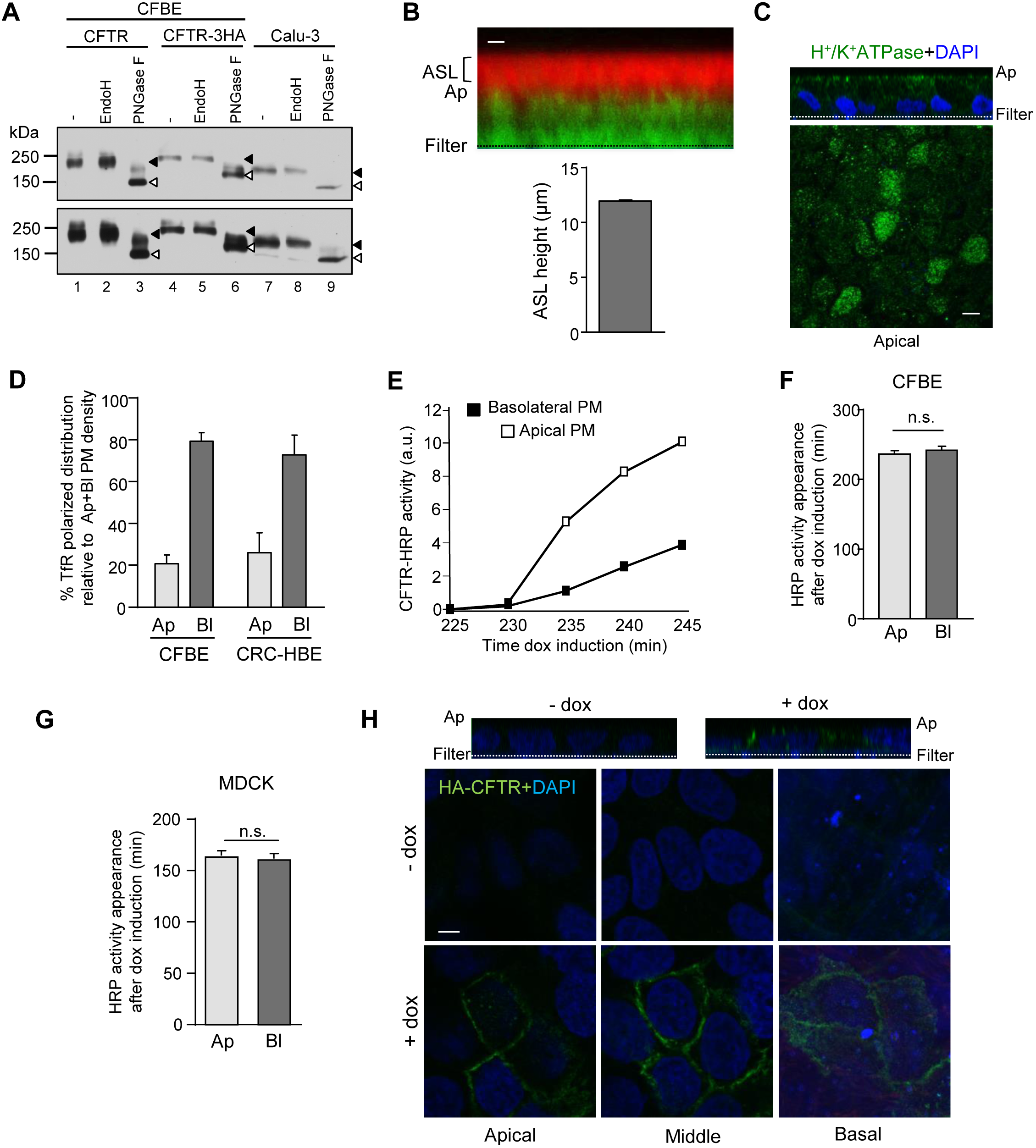
Biochemical and morphological characterization of immortalized (CFBE) and primary conditionally reprogrammed (CR-HBE) human bronchial epithelia. (A) Electrophoretic mobility of endogenous and exogenous CFTR before and after endoglycosidase digestion was visualized by immunoblotting. Cell lysates from endogenous or 3HA-tagged CFTR expressing polarized Calu-3 or CFBE cells were probed with anti-CFTR (L12B4 and M3A7) Abs before or after endoglycosidase H (endoH) or Peptide-N-Glycosidase F (PNGase F) digestion. Complex-glycosylated and deglycosylated forms of CFTR are indicated by filled and empty arrowheads, respectively. (B) ALI differentiated CR-HBE and the airway surface liquid (ASL) were stained with Cell tracker (green) and rhodamin-dextran (red), respectively (upper panel). ASL height was determined in 3-10 areas on 2 filters in 2 independent experiments as described in Methods. Bar: 10μm. (C) Differentiated CR-HBE cells at ALI interface were immunostained for H^+^/K^+^-ATPase (green) and the nuclei were visualized with DAPI (blue). Bar: 5μm. (D) Steady-state PM expression of TfR in filter-grown CFBE and CR-HBE was measured after HRP-Tf loading for 1-3 h at 37°C from the apical or basolateral compartment. HRP activity was measured at both apical and basolateral PM. Surface density is expressed as the % of total PM expression (n=4-7). (E-G) Newly synthesized CFTR-HRP concurrently arrives at both PM domains. CFTR-HRP activity was monitored after dox-induction at indicated times in the apical and basolateral compartments. Representative experiment in CFBE is shown in (E) and the mean delay of CFTR-HRP arrival time after dox induction is depicted in CFBE (n=8) and MDCK cells in (G) (n=5). (H) Specificity of basolateral CFTR staining on filter grown CFBE cells. CFBE were induced (+dox) or not (-dox) in the presence of doxycycline and incubated with anti-HA Ab from the basolateral compartment for 45 min at 37°C, while the apical compartment was supplemented with goat anti-mouse Fab. Indirect immunostaining was done on permeabilized cells and labeled CFTR was visualized by LCFM. Upper panels are representative z-sections. CFTR (green), actin (red) and DAPI (blue). Scale bar, 5μm. Data are means ± SEM on each panel. n.s. non-significant.

**Supplementary Figure 2.**
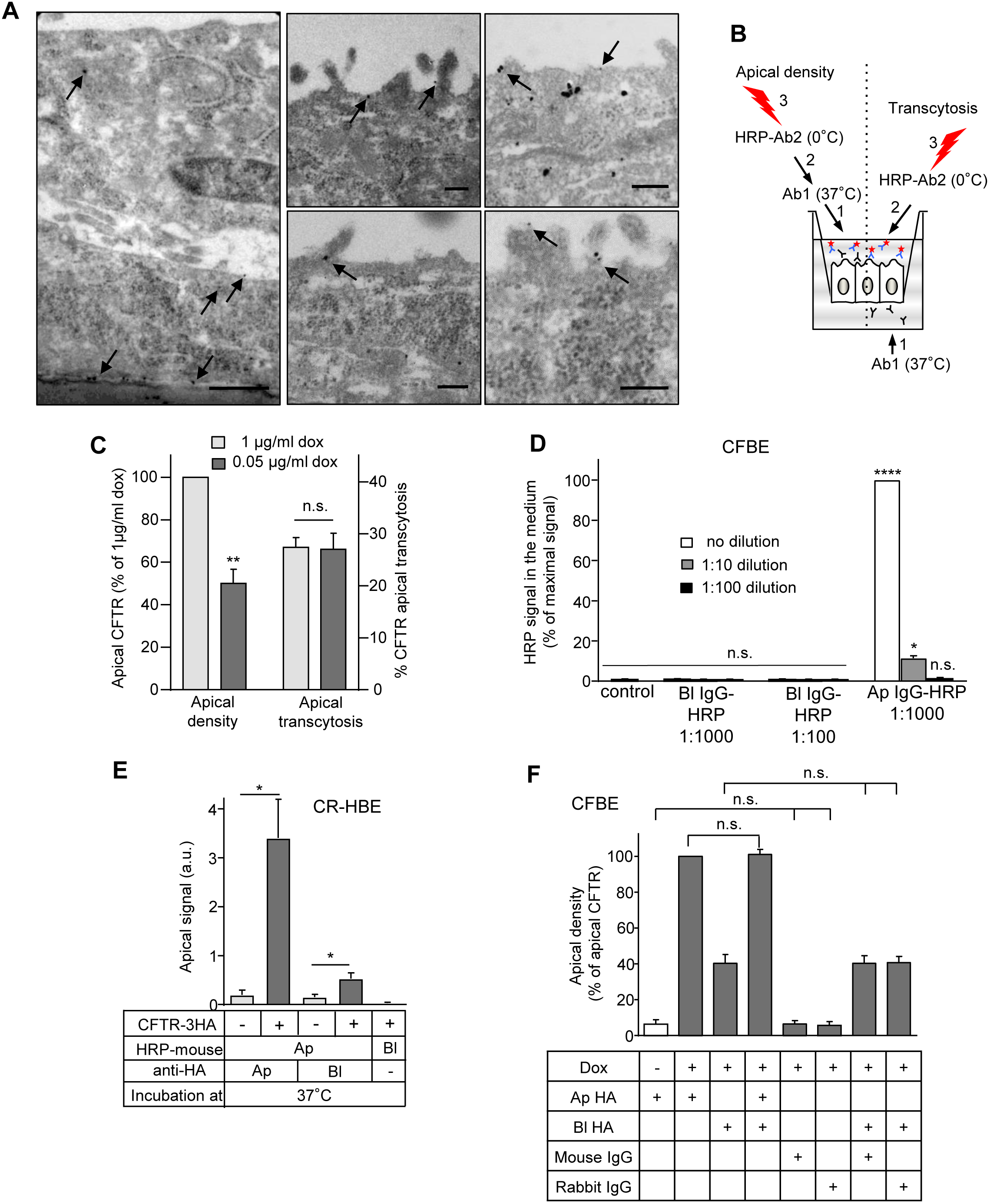
Assay development to monitor CFTR transcytosis in polarized human airway cells. (A) Immuno-EM detection of CFTR-3HA after basolateral anti-HA labeling (3h, 37°C) in CFBE. CFTR-3HA-anti-HA complexes were revealed with nanogold-conjugated anti-mouse Ab (arrows). Scale bar, 0.2 μm. (B) Apical transcytosis assay. Basolateral PM proteins were labeled with primary Ab (Ab1) at 37°C (1, right) and the Ab apical appearance was quantified by HRP-conjugated secondary Ab binding (Ab2) in the presence of Amplex Red substrate (2-3, right). In parallel samples, the steady-state CFTR density was determined at the apical PM by PM-ELISA at 37°C (1-3, left). (C) Relative transcytosis flux of CFTR is unaltered after 50% reduction of the apical CFTR expression. CFTR expression was induced in the presence of 0.05 μg/ml (light gray) or 1 μg/ml (dark grey) doxycycline. At 0.05 μg/ml doxycycline CFTR-3HA expression was ~25% of that detected in Calu-3 cells by quantitative immunoblotting (see also Fig.1A). Apical density and relative transcytosis of CFTR were measured by PM-ELISA (n=3-4). (D-F) Specificity of the PM-ELISA-based CFTR transcytosis assay in CFBE (D, F) and in CR-HBE (E) cells. (D) HRP-conjugated anti-mouse IgG were incubated in the apical (Ap) or basolateral (Bl) compartment of CFBE cells and HRP activity of diluted apical medium samples was measured in the presence of Amplex Red. Signal was normalized to the maximum HRP activity, obtained with apical IgG-HRP and statistical difference was calculated relative to untreated cells (n=3). HRP-anti-mouse Ab did not traverse the CFBE monolayer to the apical compartment. (E) No significant signal was detected at the apical PM after basolateral (3h, 37°C) incubation of HRP-anti-mouse Ab (n=2-4) in CR-HBE. (F) Similar HRP-anti-mouse IgG activity was measured at the Ap PM after Ap or Ap+Bl anti-HA incubation. No significant HRP signal was detectable at the apical PM after basolateral mouse or rabbit IgG alone or in the absence of CFTR expression with anti-HA incubation (n=3). Data are means ± SEM on each panel. n.s. non-significant, * p<0.05, ** p<0.01, **** p<0.0001.

**Supplementary Figure 3.**
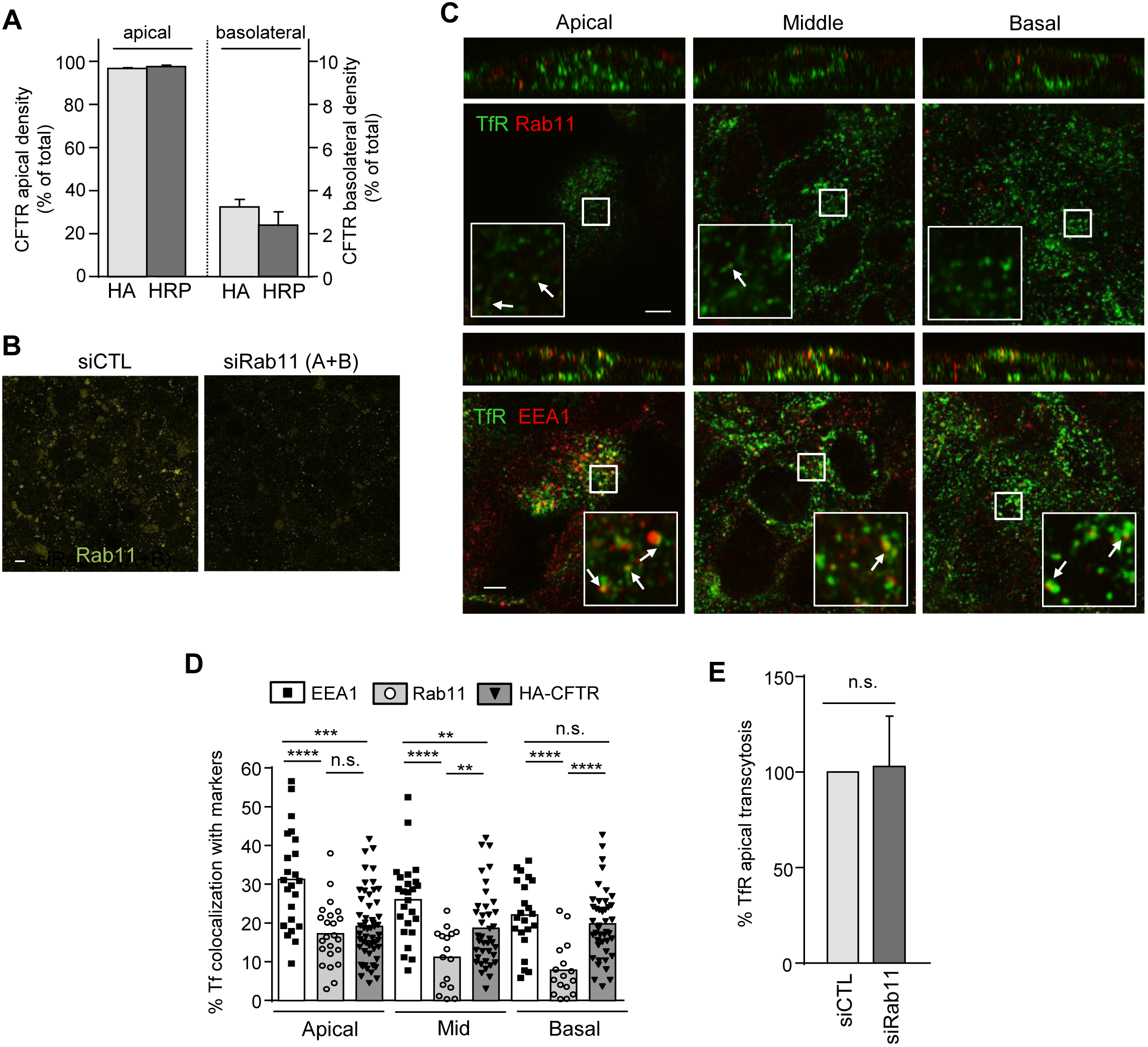
CFTR polarity and characterization of TfR transcytotic pathway in CFBE. (A) The steady-state apical and basolateral PM density of CFTR-3HA and CFTR-HRP was revealed by PM-ELISA and Amplex Red alone, respectively. CFTR domain-specific expression is indicated as percentage of its total PM density (n=3-12). (B) Specificity of Rab11 immunostaining was confirmed by ablating its expression with Rab11A+B siRNAs. Scale bar: 5 μm. (C-D) Transcytotic TfR colocalization with CFTR, EEA1 and Rab11. (C) After endogenous Tf depletion (1h, serum-free medium), Tf-Cy3 was loaded for 45min-1h at 37°C in serum-free medium (green), then cells were fixed and co-stained for EEA1 or Rab11 (red). Inserts are selected areas at ~3.5-fold increased magnification. (D) Quantitative colocalization of transcytotic TfR with EEA1, Rab11 and HA-CFTR was measured by the Mander’s coefficient. Each dot represents the Mander’s value of 16-54 R.O.I.s at the apical, middle and basal plane of CFBE, obtained from 3 independent experiments, two-tailed unpaired t-test. (E) TfR apical transcytosis was measured after 3h of Tf-HRP uptake from the basolateral compartment in CFBE cells transfected with control (CTL) or Rab11A+B siRNAs (n=3). Scale bar: 5 μm. Data are means ± SEM on each panel. n.s., non-significant, **p<0.01, ***p<0.001, **** p<0.0001

**Supplementary Figure 4.**
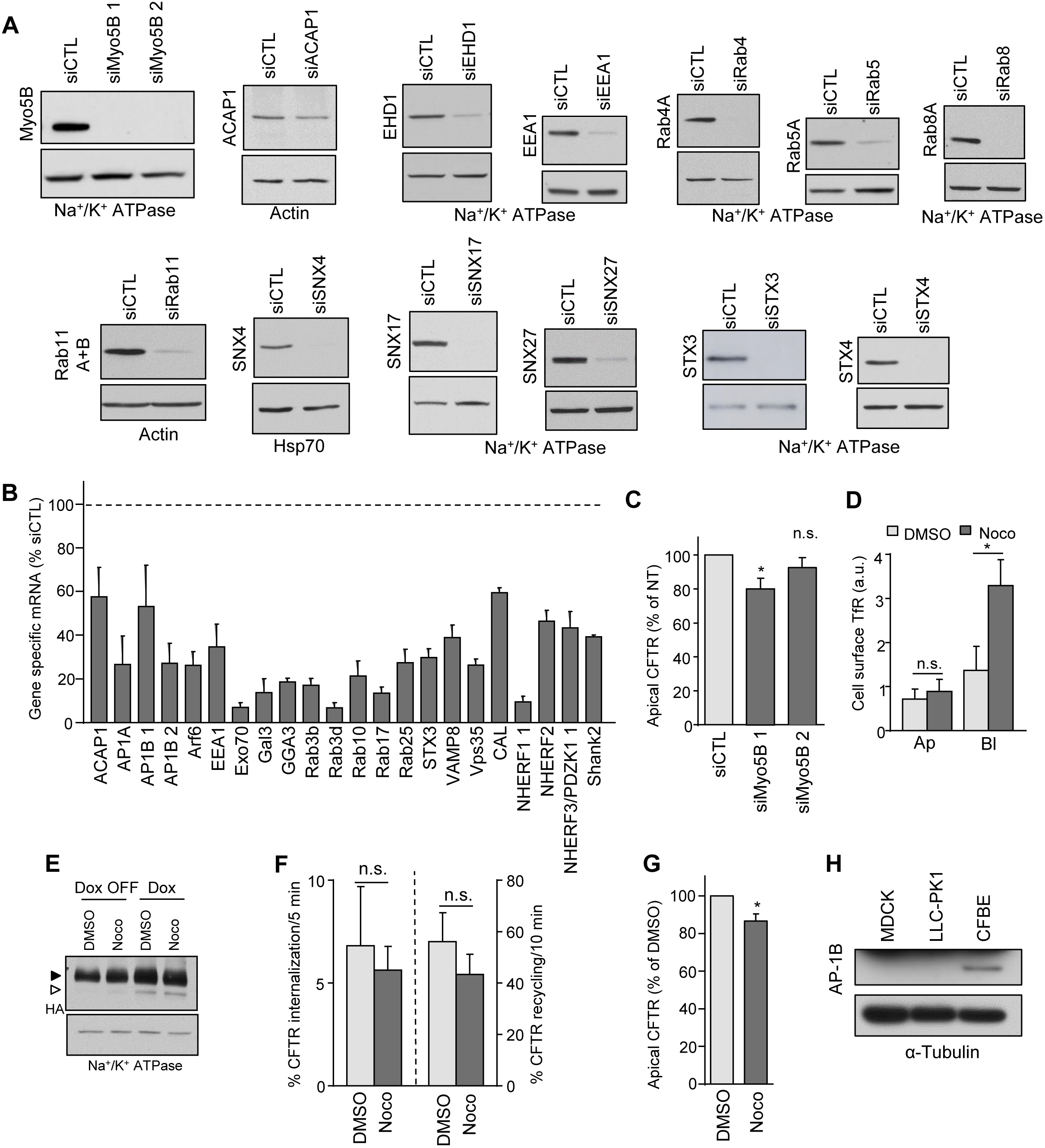
Modulators of CFTR transcytosis. (A-B) Validation of siRNA mediated silencing of target genes by immunoblotting (A) or qPCR (B) as described in Methods. Na^+^/K^+^ ATPase, β-actin and Hsp70 were used as loading controls for the immunoblots. The bar plot includes the means ± SEM of parallel samples from 3-6 independent experiments and expressed as percentage of mRNA expression of siCTL transfected CFBE cells. (C) One of the Myo5B siRNA modestly reduces CFTR apical density as compared to that of non-targeted CTL siRNA (n=4-7). (D) The apical and basolateral PM density of TfR was assayed by PM-ELISA after DMSO (1:1000) or nocodazole (Noco, 33 μM) treatment (3,5h). Ap: apical; Bl: basolateral (n=4). (E) Inhibition of transcription/translation of CFTR in dox-OFF CFBE cells decreases the core- and complex-glycosylated CFTR, detected by immunoblotting. Nocodazole (33 μM for 3.5 h) has no effect on CFTR expression relative to DMSO. Equal amounts of cell lysates were loaded. Na^+^/K^+^-ATPase was the loading control. (F) MT disruption by nocodazole (Noco, 33μM) does not significantly affect CFTR endocytosis and recycling (n=4). Internalization and recycling were measured by PM-ELISA as described in Methods. (G) The apical PM density of CFTR was assayed by PM-ELISA after DMSO (1:1000) or nocodazole (Noco, 33 μM) treatment (3,5h) (n=4). (H) Detection of AP-1B expression by immunoblotting in the indicated cell lines. 30 μg cell lysates were loaded and AP-1B expression was probed. Tubulin was used as loading control. Data are means ± SEM on each panel, n.s. non-significant, * p<0.05.

**Supplementary Figure 5.**
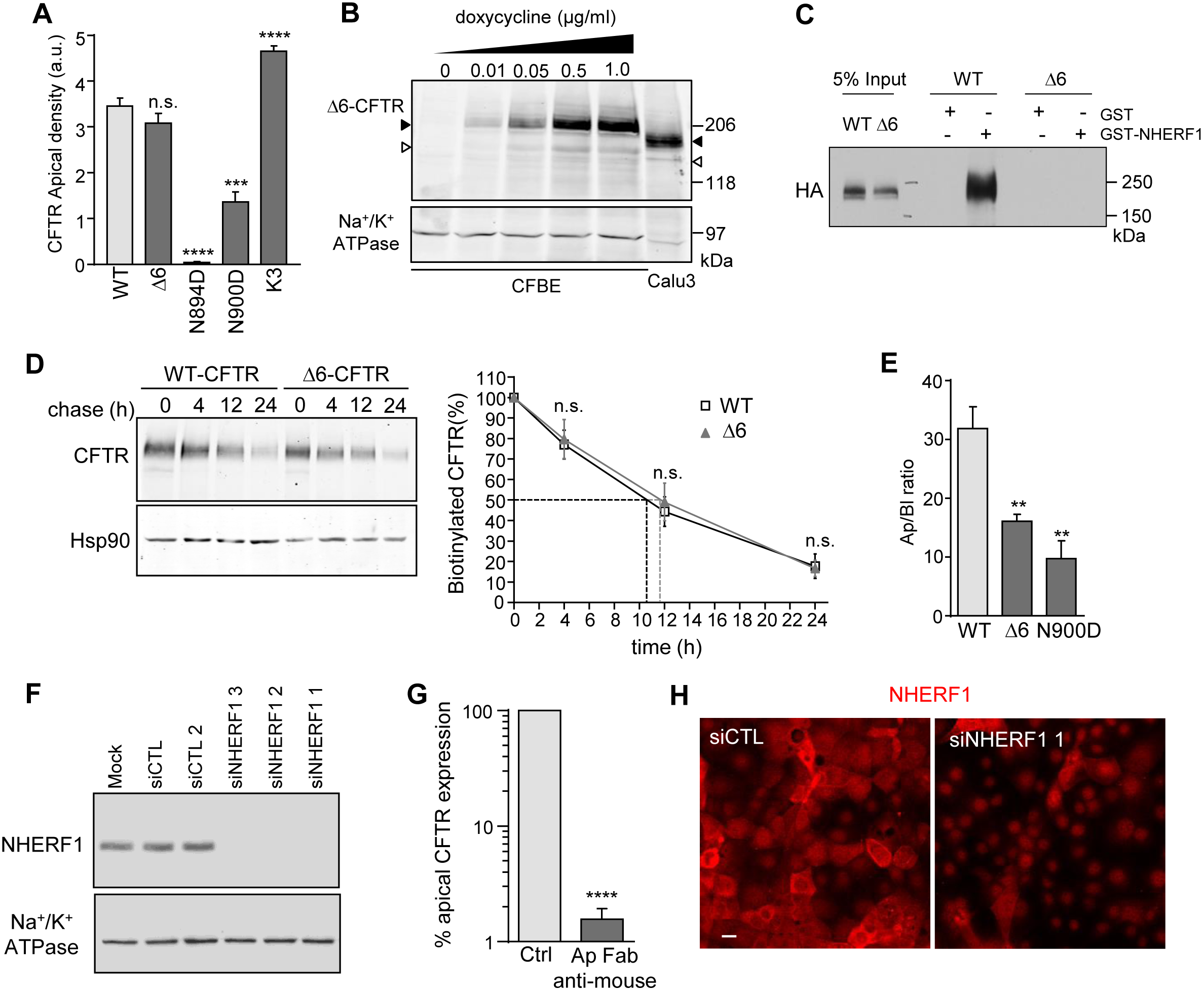
Disruption of NHERF1-CFTR interaction accelerates CFTR transcytosis. (A) The effect of putative apical or basolateral sorting signals on CFTR PM expression at the apical domain, measured by PM-ELISA. The abbreviations of the mutations are depicted in Figure 6A (n=3-40, two-tailed unpaired t-test). (B) Δ6-CFTR was induced with doxycycline at the indicate concentration for 3 days. CFTR-3HA expression was compared to that of endogenous CFTR of Calu-3 cells in equal amounts of cell lysates by immunoblotting with anti-CFTR Abs. Na^+^/K^+^-ATPase was used as loading control. (C) WT but not Δ6-CFTR binds to recombinant NHERF1. Solubilized WT and Δ6-CFTR were pulled down with recombinant GST- or GST-NHERF1, immobilized on glutathione Sepharose beads. CFTR was probed by immunoblotting together with 5% input of cell lysates with anti-HA Ab and ECL. Representative of 3 independent experiments. (D) Cell surface stability of WT- and Δ6-CFTR. The PM turnover of CFTR variants was measured by biotinylation with NHS-SS-biotin at 0°C, followed by the indicated chase at 37°C. Biotinylated proteins were immunoprecipitated on monomeric avidin beads and CFTR was visualized by quantitative immunoblotting. Hsp90 was used as loading control. Right panel: quantification of CFTR remaining during the chase (n=3). (E) WT-, Δ6-, and N900D-CFTR expression ratios at the Ap and Bl (Ap/Bl) PM were measured by comparing the signal obtained at both domains by PM ELISA (n=4-15, two-tailed unpaired t-test). (F) NHERF1 siRNA silencing efficiency was monitored by immunoblotting. Na^+^/K^+^ ATPase was used as loading control. (G) The blocking efficiency of anti-HA Ab by goat anti-mouse Fab. After anti-HA labeling of CFTR at the apical PM, CFBE cells were incubated on ice with PBSCM (control) or PBSCM-BSA0.5% + goat anti-mouse Fab (1:75). Remaining anti-HA Ab was probed with HRP-conjugated anti-mouse Ab on ice and the specific HRP activity was quantified in the presence of Amplex Red (n=8). (H) Specificity of NHERF1 detection was confirmed by immunostaining NHERF1 in control (CTL) or NHERF1_1 siRNA transfected CFBE cells. Scale bar: 20μm. Data are means ± SEM on each panel. n.s. non-significant, * p<0.05, ** p<0.01, *** p<0.001, **** p<0.0001.

**Supplementary Figure 6.**
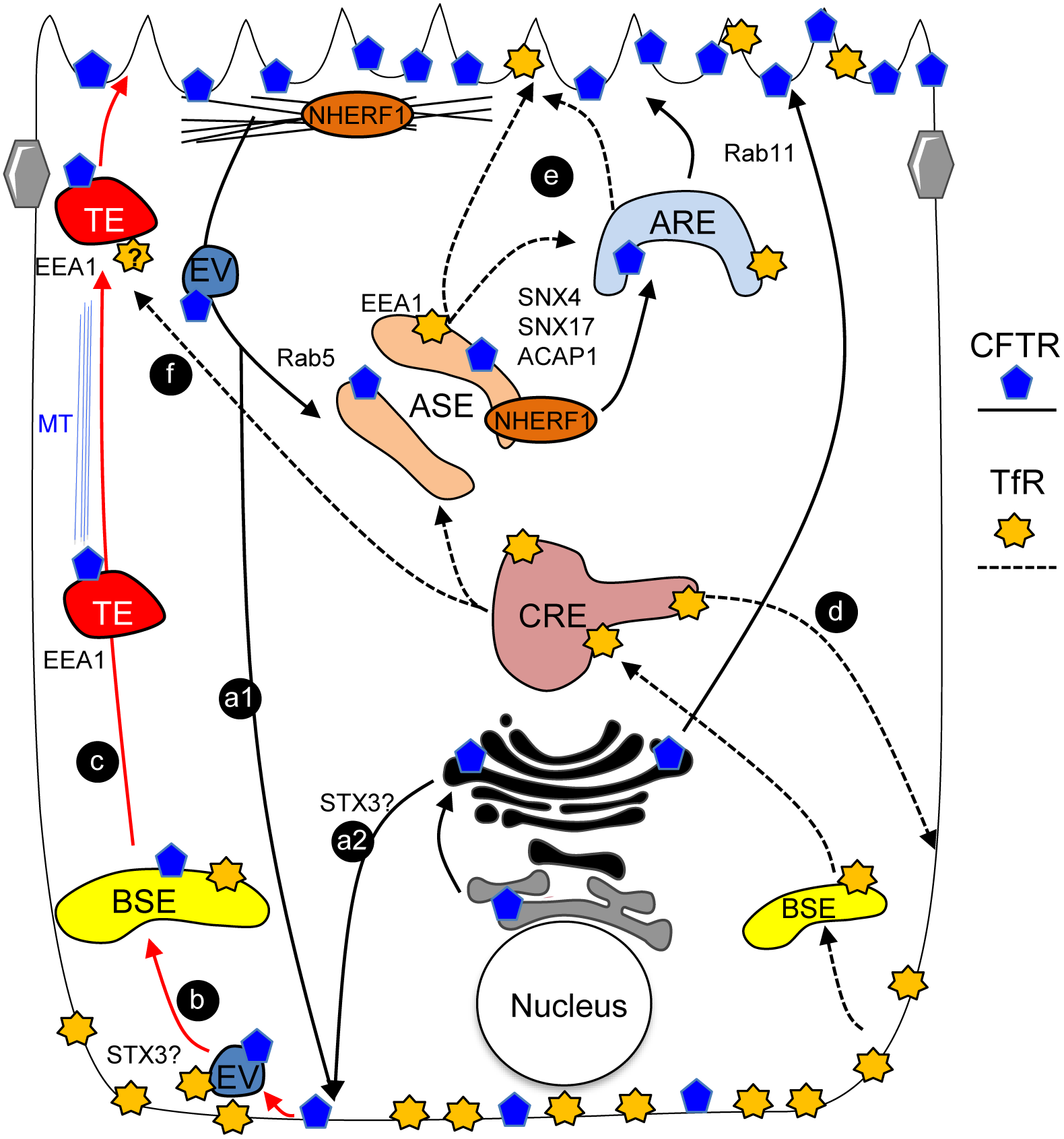
Working model of CFTR transcytosis in airway epithelia. CFTR basolateral missorting occurs from apical endosomes (a1) and at the TGN along the biosynthetic pathway (a2). (b) Missorted basolateral CFTR is retrieved from the basolateral PM into basolateral sorting endosomes (BSE). (c) CFTR is packaged into EEA1 positive transcytotic endosomes (TE) from BSE and ferried to the apical pole in a microtubules-dependent pathway. (d) The basolateral internalization, recycling and (e) transcytotic route of TfR is indicated by dashed lines for comparison (Perez Bay et al., 2013). While in AP1B-KD MDCK cells transcytotic TfR traverses the ARE, in CFBE cells the ARE may have a minor role in TfR delivery to the apical PM. CFTR and TfR may also use overlapping apical exocytic platform (f). ASE, Apical Sorting Endosome; ARE, Apical Recycling Endosome; BSE: Basolateral Sorting Endosome; CRE: Common Recycling Endosome; EV, Endocytic Vesicles; Bundled blue lines, microtubules (MT); interlaced black lines: cortical F-actin.

**Table S1.**
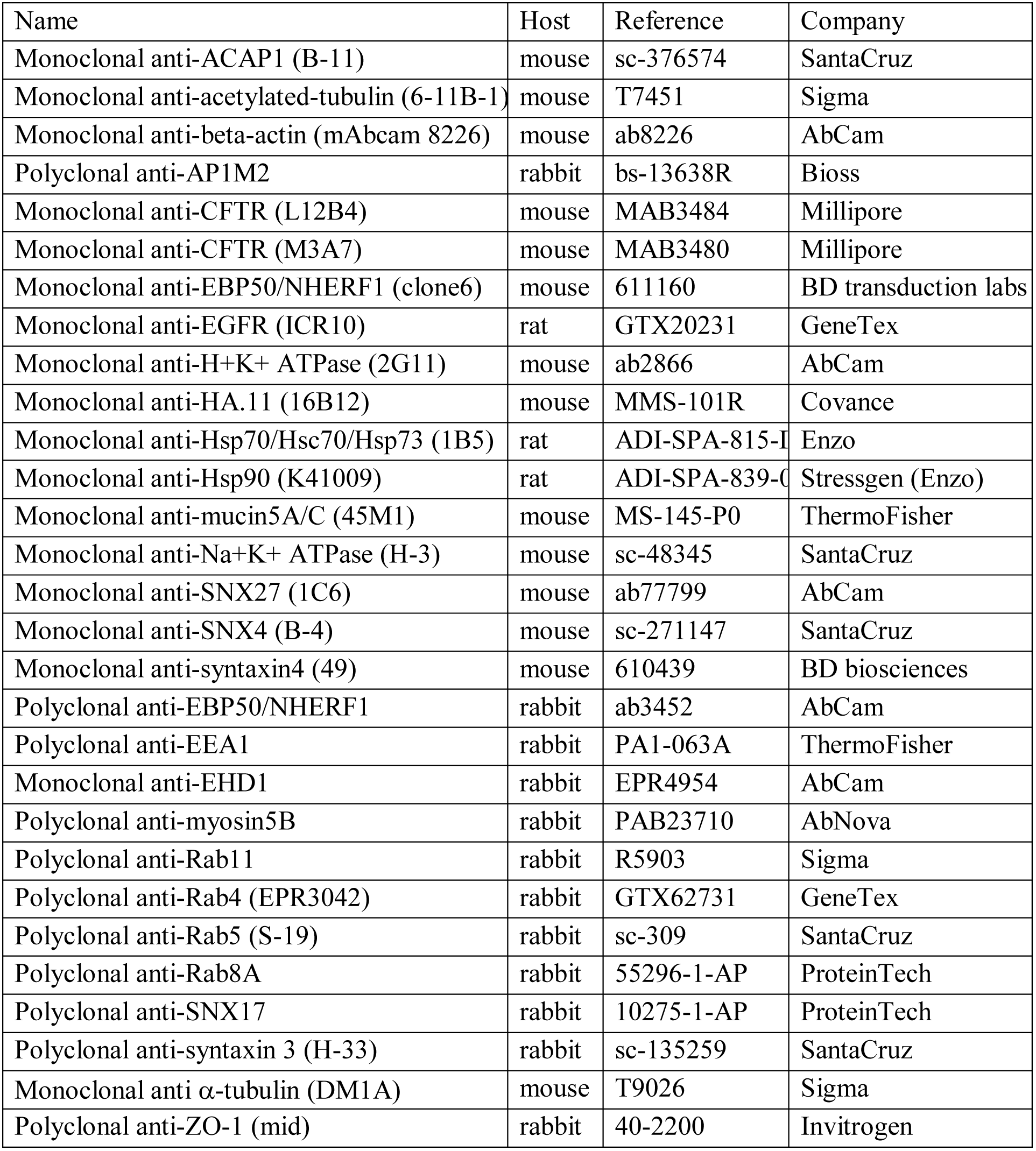
Antibodies used in this study.

**Table S2.** siRNAs used in this study. Normal and bold letters indicate predesigned Qiagen/Origene siRNAs and Invitrogen stealth RNAi siRNAs, respectively.

| Gene | Product name |
| --- | --- |
| CTL | **Control stealth Med GC** |
| NHERF1 1 | **HSS145123** |
| NHERF1 2 | Hs_SLC9A3R1_5 |
| NHERF2 | **HSS113866 ; HSS113867** |
| PDZK1 1 | Hs_PDZK1_8 |
| PDZK1 2 | **HSS107800 ; HSS182218** |
| Shank2 1 | **HSS117944 ; HSS177049** |
| Shank2 2 | Hs_SHANK2_9 ; Hs_SHANK2_10 |
| CAL | **HSS126013** |
| NHERF4 | **HSS188224** |
| Rab3b | Hs_RAB3B_3 ; Hs_RAB3B_4 |
| Rab3d | Hs_RAB3D_3 ; Hs_RAB3D_4 |
| Rab4 | Hs_RAB4A_5 ; Hs_RAB4A_6 |
| Rab5 | Hs_RAB5A_5 ; Hs_RAB5A_8 |
| Rab8 | Hs_RAB8A_4 ; Hs_RAB8A_5 |
| Rab10 | Hs_RAB10_3 ; Hs_RAB10_5 |
| Rab11 | Hs_RAB11A_5 ; Hs_RAB11A_6 ; Hs_RAB11B_1 ; Hs_RAB11B_2 |
| Rab17 | Hs_RAB17_4 ; Hs_RAB17_5 |
| EEA1 | Hs_EEA1_5 ; Hs_EEA1_7 ; Hs_EEA1_9 |
| Rab25 | Hs_RAB25_1 ; Hs_RAB25_3 |
| EHD1 | Hs_EHD1_3 |
| Vps35 | Hs_VPS35_5 ; Hs_VPS35_6 ; Hs_VPS35_8 |
| ACAP1 | Hs_CENTB1_1 ; Hs_CENTB1_5 |
| Arf6 | Hs_ARF6_5 ; Hs_ARF6_8 |
| GGA3 | Hs_GGA3_1 ; Hs_GGA3_9 |
| SNX4 | Hs_SNX4_2 ; Hs_SNX4_7 |
| SNX17 | Hs_SNX17_1 ; Hs_SNX17_2 |
| SNX27 | Hs_SNX27_5 ; Hs_SNX27_6 |
| STX3 | SR321926-A |
| STX4 | Hs_STX4A_2 ; Hs_STX4_1 |
| Vamp8 | **HSS189495** |
| Exo70 | Hs_EXOC7_5 ; Hs_EXOC7_6 |
| Gal3 | Hs_LGALS3_7 ; Hs_LGALS3_3 ; **Hs_LGALS3_8** |
| AP1A | Hs_AP1M1_1 ; Hs_AP1M1_6 |
| AP1B 1 | Hs_AP1M2_5 ; Hs_AP1M2_6 ; Hs_AP1M2_8 |
| AP1B 2 | Hs_AP1M2_7 ; Hs_AP1M2_9 ; Hs_AP1M2_10 |
| Myo5B 1 | **HSS181415** |
| Myo5B 2 | Hs_MYO5B_7 ; Hs_MYO5B_10 |

**Table S3.**
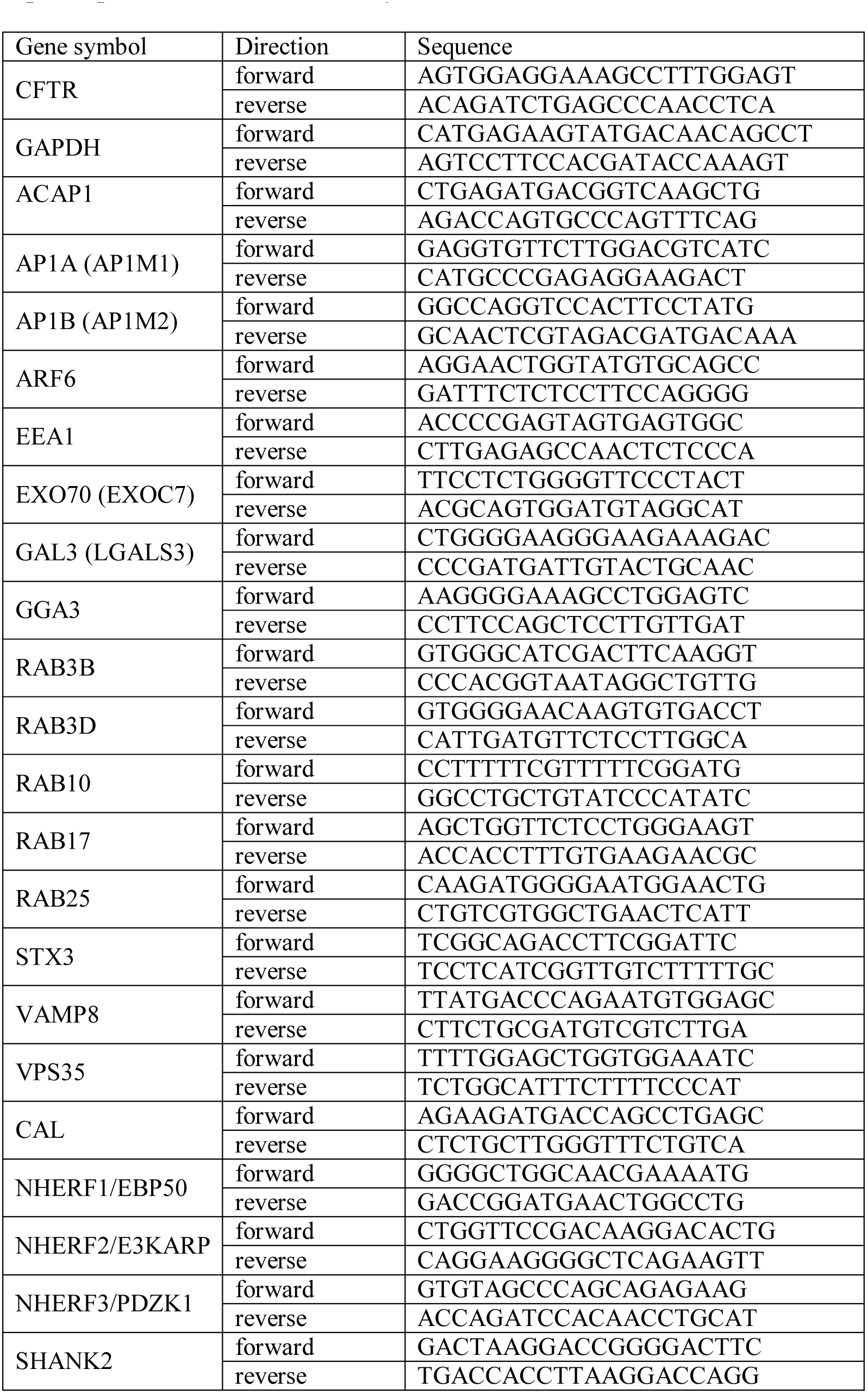
qPCR primers used in this study.

